# A gravity-based three-dimensional compass in the mouse brain

**DOI:** 10.1101/570382

**Authors:** Dora E Angelaki, J Ng, AM Abrego, HX Cham, JD Dickman, J Laurens

## Abstract

Head direction cells in the mammalian limbic system are thought to function as an allocentric neuronal compass. Although traditional views hold that the compass of ground-dwelling species is planar, we show that head-direction cells in the rodent thalamus, retrosplenial cortex and cingulum fiber bundle are tuned to conjunctive combinations of azimuth, pitch or roll, similarly to presubicular cells in flying bats. Pitch and roll orientation tuning is ubiquitous, anchored to gravity, and independent of visual landmarks. When head tilts, azimuth tuning is affixed to the head-horizontal plane, but also uses gravity to remain anchored to the terrestrial allocentric world. These findings suggest that gravity defines all three degrees of freedom of the allocentric orientation compass, and only the azimuth component can flexibly remap to local cues in different environments. Collectively, these results demonstrate that a three-dimensional, gravity-based, neural compass is likely a ubiquitous property of mammalian species, including ground-dwelling animals.

## Introduction

Head direction (HD) cells, which encode allocentric head orientation analogous to a "neural compass" are a primordial component of the mammalian (Taube 2007; Finkelstein et al. 2015) and fly (Seelig and Jayaraman, 2015; Green et al., 2017; Turner-Evans et al., 2017; Kim et al., 2017) spatial navigation system. Originally identified in the rat dorsal presubiculum (Taube et al. 1990a,b; Preston-Ferrer et al. 2016; Simonnet et al. 2018), HD cells have been subsequently found in multiple brain regions, including the anterior thalamus (Taube et al. 1995; Peyrache et al. 2015, 2017; Page et al. 2017; Shinder and Taube 2011, 2014, 2019), parasubiculum (Kornienko et al. 2018), retrosplenial cortex (Chen et al. 1994a,b; Cho et al. 2001; Jacob et al. 2017; Lozano et al. 2017), and entorhinal cortex (Sargolini et al. 2006; Kornienko et al. 2018; Park et al. 2018).

HD cells have been traditionally known for encoding head orientation in the horizontal plane (azimuth) by firing when the animal faces a particular direction. Yet, HD cells in the bat presubiculum were recently shown to respond in vertical planes, i.e. when the head tilts in pitch or roll (Finkelstein et al. 2015). Gravity is a ubiquitous cue for allocentric vertical and may thus provide an Earth constant reference for the brain compass. However, Finkelstein et al. (2015) did not investigate whether tilt tuning is anchored to gravity. In parallel, Laurens et al. (2016) identified gravity-anchored tilt tuning in the anterior thalamus of rhesus macaques, but did not test whether gravity-tuned cells are traditional azimuth-tuned HD cells. Thus, the two observations (gravity-anchored tilt tuning in macaques and 3D HD tuning in bats) have remained segregated. Furthermore, tuning to tilt has never been shown in rodents, and some researchers proposed that it may be restricted to aerial and tree-dwelling species but absent in ground-dwelling species like rodents (Calton and Taube 2005; Shinder and Taube 2019).

Here we test the hypothesis that gravity-anchored tilt signals and visually-anchored azimuth signals converge onto rodent HD cells to yield a sense of 3D head orientation (Laurens and Angelaki, 2018; **Fig. 1a**). As in bat presubiculum, we first show that HD cells in the mouse anterior thalamus, retrosplenial cortex and cingulum are tuned to combinations of azimuth and tilt. We then show that, not only does gravity anchor tilt tuning, but it also participates in updating azimuth to preserve allocentric invariance during 3D motion (Laurens and Angelaki, 2018). Finally, we also show that gravity-anchored tilt (vertical) tuning and gravity-defined azimuth tuning interact multiplicatively to endow a neural correlate of 3D spatial orientation. Thus, a 3D orientation compass is not a specialized property limited to areal species, but may instead represent a ubiquitous property throughout many chapters of animal evolution.

**Figure 1.**
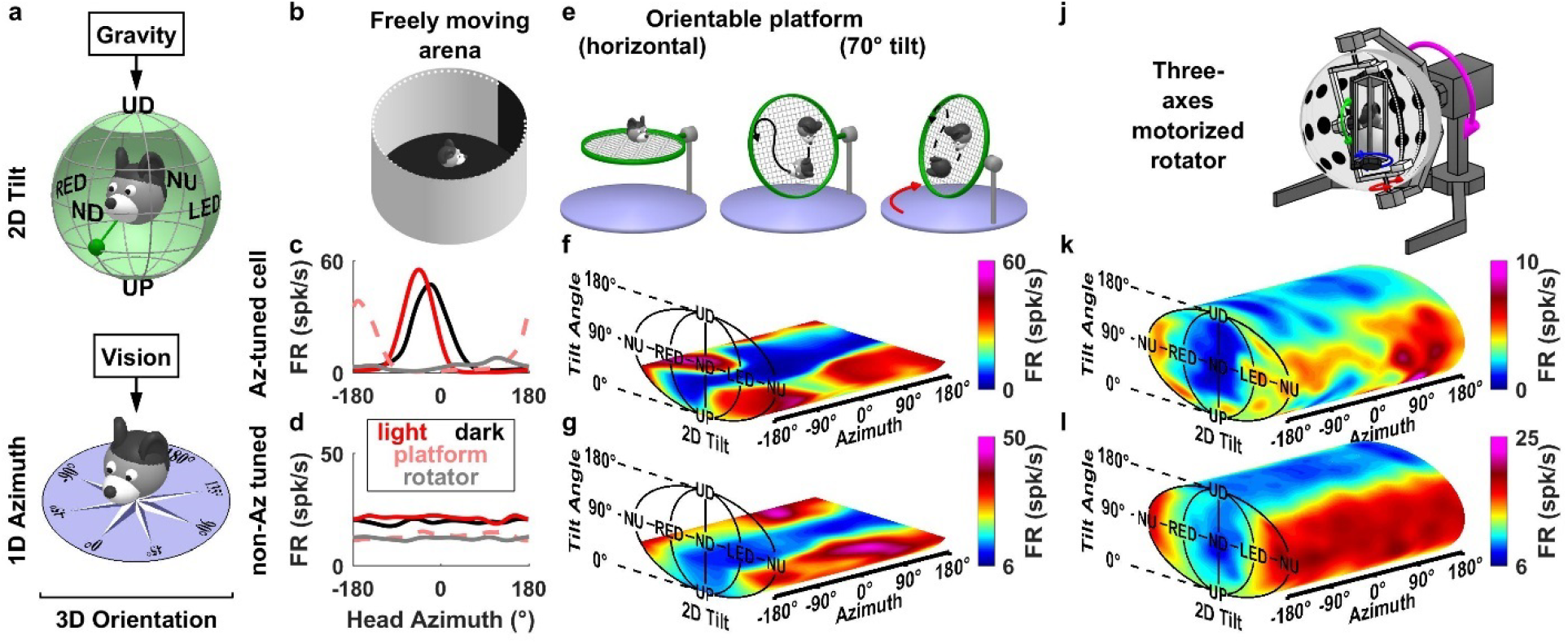
Example cells tuned to tilt. **(a)** Schematic representation of the topologies of tilt (2D, spherical - top) and azimuth (1D, circular - bottom). **(b)** Schematic of the arena used to identify azimuth-tuned cells during free foraging in the horizontal plane. **(c-d)** Example azimuth tuning of a ‘traditional HD celľ, i.e. tuned to azimuth (‘Az-tuned’) in the ADN (c) and another cell not tuned to azimuth (‘non-Az tuned’ cell) in the CIN (d), as the mouse walks freely in light (red) and darkness (black) in a horizontal arena (shown in b), on a platform oriented horizontally (shown in e, left) (broken pink lines) and in the rotator (shown in j) (gray lines). The Az-tuned cell exhibited a significant tuning with different preferred directions (PD) in all setups, although response amplitude was strongly attenuated in the rotator (compare gray with red/pink lines). **(e)** Schematic of a 3D orientable platform used to measure 3D tuning. Two-degrees-of-freedom allow the platform to be tilted and re-oriented. **(f,g)** Tuning curves for the two cells in c and d, obtained from responses as the mouse foraged on the orientable platform (shown in e). Firing rate is shown as a heat map in 3D space (see **Suppl. Movies 1,2**). Note that tuning curves are restricted within a narrow sector of ~60° around upright (see Methods). **(j)** Schematic of a ‘rotator’ used to measure complete 3D tuning curves. **(k,l)** Tuning curves for the two cells in (c,f) and (d,g) as the mouse was passively re-oriented uniformly throughout the full 3D space using the rotator (see **Suppl. Movie 3,4,5**). Note that, unlike Az-tuning, tilt PDs are maintained across different environments.

### A 3D compass in the mouse brain

We used tetrodes to record extracellularly from the antero-dorsal nucleus of the thalamus (ADN; n = 4 mice; **Suppl. Table 1**), retrosplenial cortex (RSC; n=2 mice; **Suppl. Table 1**), and cingulum fiber bundle (CIN; n = 4 mice; **Suppl. Table 1**). The CIN carries projection fibers from the ADN and RSC (Domesick 1970; van Groen and Vyss 1990; 1995; Bubb et al. 2018). Cells were exclusively selected based on spike isolation (**Suppl. Fig. 1**) and recording locations were verified post-mortem (**Suppl. Fig. 2**). Based on their head orientation during free foraging in a horizontal arena (**Fig. 1b**; summarized in **Suppl. Fig. 3**), cells were characterized as azimuth (Az)-tuned (i.e. ‘traditional’ HD) cells in light (**Fig. 1c**, red) and darkness (see example in **Fig. 1c**, black) or azimuth-untuned (see example in **Fig. 1d**).

Neurons were then characterized as animals walked on a 3D orientable platform (**Fig. 1e**) that could be re-positioned so that animals foraged through multiple tilt angles ranging between 0° (upright) and ~60° (**Suppl. Fig. 4**). We computed tilt tuning curves in spherical coordinates, with 2 degrees of freedom: tilt angle from upright: α (range: 0-180°), and tilt orientation: γ (range: 0-360°; see **Suppl. Fig. 5** for definitions of right/left ear-down [RED/LED] and nose-up [NU]/down [ND] orientations). For simplicity of illustration, 2D tilt tuning curves are shown in a planar representation using an equal-area Mollweide projection, and the 1D azimuth axis is unfolded. Next, the neurons’ tuning was plotted as a color map in a volume formed by this tilt plane and the azimuth axis (**Fig. 1f,g; Suppl. Movie 1**). Both example neurons in **Fig. 1** exhibited tilt tuning in the restricted 3D orientation space, characterized by an increased firing rate at a preferred tilt (**Fig. 1f,g**, NU for both cells; see **Suppl. Movie 2**), with peak-to-trough amplitudes of 32 spk/s and 23 spk/s, respectively when averaged across all azimuth angles. In addition, with the platform in the earth-horizontal orientation, the cell classification as Az-tuned or Az-untuned persisted (**Fig. 1c,d**, dashed pink curves).

Next, animals were transferred to a multi-axis rotator (**Fig. 1j**) that sampled 3D orientation uniformly to measure the neurons’ full 3D tuning curves (**Suppl. Fig. 6; Suppl. Movie 3**). The example azimuth-tuned cell in **Fig. 1c, f** maintained both azimuth (**Fig. 1c**, gray) and tilt (**Fig. 1k**; see also **Suppl. Movie 4**) tuning in the rotator, although its peak firing rate had decreased from ~60 spikes/s to ~10 spikes/s. While the example cell’s preferred direction (PD) in azimuth differed across environments (**Fig. 1c**, compare solid red, dashed pink and gray lines, with PDs at −46°, −163°, and 114°, respectively), its tilt PD was relatively constant at NU (**Fig. 1k**: [α=83°, γ=180°], as compared to **Fig. 1f**: [α=56°, γ=130°; note that tuning was not sampled at higher tilt angles]). Similarly, the example Az-untuned cell exhibited a tilt PD at [α=97°, γ=168°] (**Fig. 1l; Suppl. Movie 5**), as compared to [α=48°, γ=150°] (**Fig. 1g**). Thus, tilt tuning recorded when animals were passively re-oriented in the rotator had similar spatial properties to that when moving freely in 3D.

We used identical criteria to classify neurons as azimuth-tuned or tilt-tuned (Methods; **Suppl. Fig. 7**). When tested on the platform, we found that tilt tuning was widespread in Az-tuned (traditional HD) cells classified based on their responsiveness in the horizontal arena (of **Fig. 1b**), as summarized in **Fig. 2**. Specifically, out of 29 ADN neurons recorded on the platform, 25 (86%) were classified as azimuth-tuned (**Fig. 2a**, Venn diagram). Amongst these, 24 (96%) were also tuned to tilt and are subsequently called ***conjunctive (azimuth and tilt)*** HD cells (solid red symbols in **Fig. 2**). A sizeable population of Az-tuned cells were also recorded in RSC and CIN (49% and 38% of recorded cells, respectively). Of these, 58% (RSC) and 75% (CIN) were conjunctive cells.

**Figure 2:**
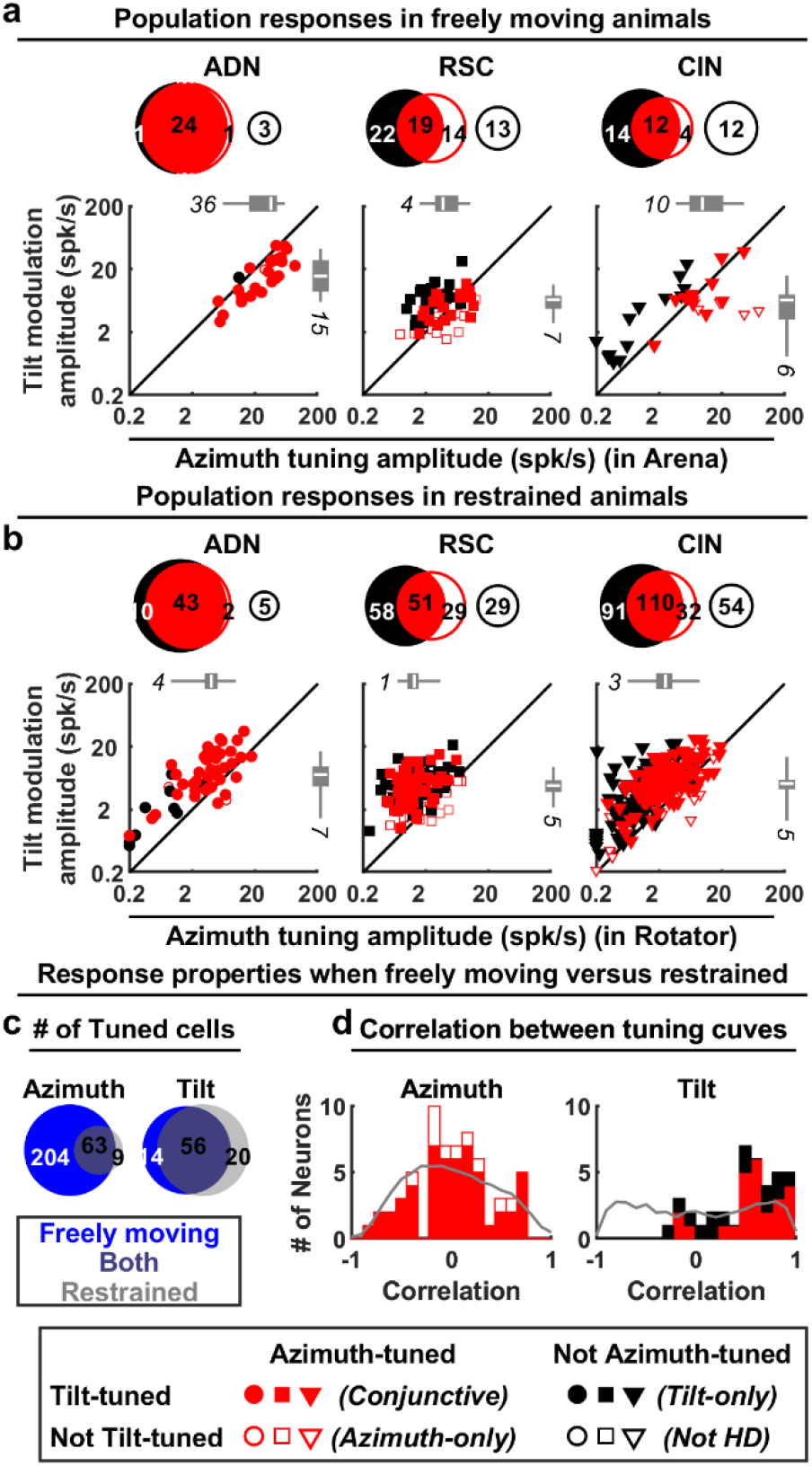
Population azimuth and tilt tuning in freely moving vs. restrained animals, segregated by area. **(a)** Summary of tuning prevalence during unrestrained motion. Azimuth tuning was derived from data in the freely moving arena (**Fig. 1b**). Tilt tuning was derived from data on the 3D platform (**Fig. 1e**). For each panel, Venn diagrams (top) indicate the number of tilt-tuned (filled black discs) and azimuth-tuned (red disks) cells. Conjunctive cells appear at the intersection of these disks. Open disks illustrate cells responsive to neither tilt nor azimuth. The scatterplots (bottom) indicate the peak-to-trough modulation amplitude of responsive cells, computed from Gaussian (tilt tuning) or von Mises (azimuth tuning) fits; see Methods. The boxes and whiskers represent the median (white line), 95% confidence interval (boxes) and upper/lower quartiles (whiskers) of the azimuth modulation of azimuth-tuned cells (top) and tilt modulation of tilt-tuned cells (right). Different symbols (based on recorded area) are color-coded based on cell type (Conjunctive: filled red; Azimuth-only: open red; Tilt-only: filled black). **(b)** Summary of prevalence of tilt tuning in restrained animals rotated passively in the rotator (**Fig. 1j**) and azimuth tuning (when moving freely). Format as in panel a. **(c)** Comparison of responsiveness for cells tested in both restrained and freely foraging animals. Venn diagrams with the number of cells tuned when moving freely (blue) in the arena (azimuth tuning) or 3D platform (tilt tuning) and restrained in the rotator (grey). Cells tuned under both conditions appear at the intersection of both discs. **(d)** Pixel-by-pixel correlation of the fitted azimuth (left) and tilt (right) tuning curves (only cells tuned under both freely moving and restrained conditions are included). For tilt tuning, the rotator data were re-analyzed by restricting tilt angles up to 60° (to match the conditions in the platform). Grey: expected distribution if tuning curves shift randomly (H0), computed by randomly shuffling the cells.

Tilt tuning on the platform was also seen in Az-untuned cells (solid black disks and symbols in **Fig. 2a**). Thus, tilt tuning was common, observed in all areas, regardless of Az-tuning, with 92/139 (66%) tilt-tuned cells. A total of 74/139 (53%) cells were Az-tuned, and tilt and azimuth tuning showed some overlap across neurons. Tilt-tuned cells were slightly (7%) more likely to be Az-tuned and reciprocally Az-tuned cells were slightly (8%) more likely to be tilt-tuned (Chi-square test, p=0.03). The peak-to-trough modulation of tilt and azimuth tuning was comparable (**Fig. 2a**; see also **Suppl. Fig. 7d-f**), although tilt tuning amplitude was lower in conjunctive ADN cells (Wilcoxon paired rank test; p=10^−4^ in ADN, p=0.4 in RSC, p=0.6 in CIN). These results indicate that tilt signals are an inherent component of the mouse HD system during natural behavior; thus, the term “HD cell” should refer to both tilt-tuned cells as well as azimuth-tuned cells. However, 3D characterization in freely moving mice was limited by the explorable space.

### Spatial properties of tilt tuning in 3D

To characterize tuning uniformly in 3D space, 514 (60 ADN, 167 RSC, 287 CIN) neurons were tested in the rotator. Seventy-one percent (363) of these cells were significantly tuned to tilt (88% ADN, 65% RSC and 70% CIN). Similar to responses obtained to foraging on the platform, tilt-tuned cells were slightly (4%) more likely to be Az-tuned and reciprocally Az-tuned cells were slightly (6%) more likely to be tilt-tuned (Chi-square test, p=0.003).

In line with previous studies (Shinder and Taube, 2011; 2014) and the example cell in **Fig. 1c,k**, tuning modulation amplitude was attenuated when animals were restrained in the rotator (scatter plots in **Fig. 2b** vs. **Fig. 2a**). Azimuth tuning amplitude was attenuated by a factor of 4.9 (**Suppl. Fig. 8a**). As a result, only a minority of neurons tuned to azimuth when moving freely were significantly tuned to azimuth when restrained in the rotator (63/267; **Fig. 2c**, left panel). Nevertheless, the PDs of multiple azimuth-tuned HD cells had consistent angles relative to each other when moving freely and in the rotator (**Suppl. Fig. 8d**), suggesting that an attractor network structure of the population of azimuth-tuned cells was maintained in the rotator. Thus, other than the smaller response magnitude, azimuth (and, as will be shown below, tilt) responses measured in the rotator are representative of the neurons’ natural responses.

Tilt tuning magnitude was also attenuated in the rotator, although to a lesser extent (by a factor of 2.4; **Suppl. Fig. 8b**). A minority of cells tested both on the orientable platform and the rotator were tilt tuned only during stimulation in the rotator because of the larger sampling of 3D space as compared to foraging on the platform that was limited to 60° tilt. In contrast, some cells were significantly tuned only when moving freely because the response magnitudes were larger (**Suppl. Fig. 8b**). Nevertheless, a large proportion of cells (62%) were tuned to tilt under both conditions (**Fig. 2c**, right panel), indicating that tilt tuning is conserved across free locomotion and restrained, passive motion conditions.

We then compared tuning curves in freely moving and restrained animals by computing their pixel-by-pixel correlations. This revealed an important difference between azimuth and tilt tuning. Because azimuth curves shifted randomly between environments (**Suppl. Fig. 8c**), their pixel-by-pixel correlations were uniformly distributed (median=0.07; Kolmogorov-Smirnov test p=0.15; **Fig. 2d**, left). In contrast, tilt tuning was preserved (median=0.58; p<10^−5^; **Fig. 2d**, right), as expected if the tilt compass was anchored to a common reference: gravity (**Fig. 1a**; see direct testing of this hypothesis below).

The majority of 3D tilt tuning curves were unimodal (as in the example responses of **Fig. 1k,l**, see also **Suppl. Fig. 9a**). However, bimodal (**Suppl. Fig. 9b**) or complex-shaped (**Suppl. Fig. 9c**) tuning curves were also encountered. Regardless of their shape, tilt tuning curves were well fitted with Gaussian functions (**Suppl. Fig. 9,10**), which allowed quantification of tilt preferred directions in 3D. As illustrated in **Fig. 3a**, PDs were scattered across the full range of angles, suggesting that the population of tilt-tuned neurons could provide a code for tilt orientation.

Yet, the distribution was not uniform, with an overrepresentation of ND pitch PDs and an underrepresentation of PDs in the roll plane (LED/RED, grey sectors) ADN: p<10^−6^; RSC: p<2.10^−3^; CIN: p<10^−10^; Chi Square). In contrast, there was no significant difference in PDs between the upper vs. lower hemispheres of the 3D space (i.e. tilt angles lower or higher than 90° (p>0.1 in all areas, Chi Square). These results, showing a dominance of pitch-tuned over roll-tuned cells, are consistent with those previously described for bats (Finkestein et al., 2015) and monkeys (Laurens et al., 2016).

**Fig 3:**
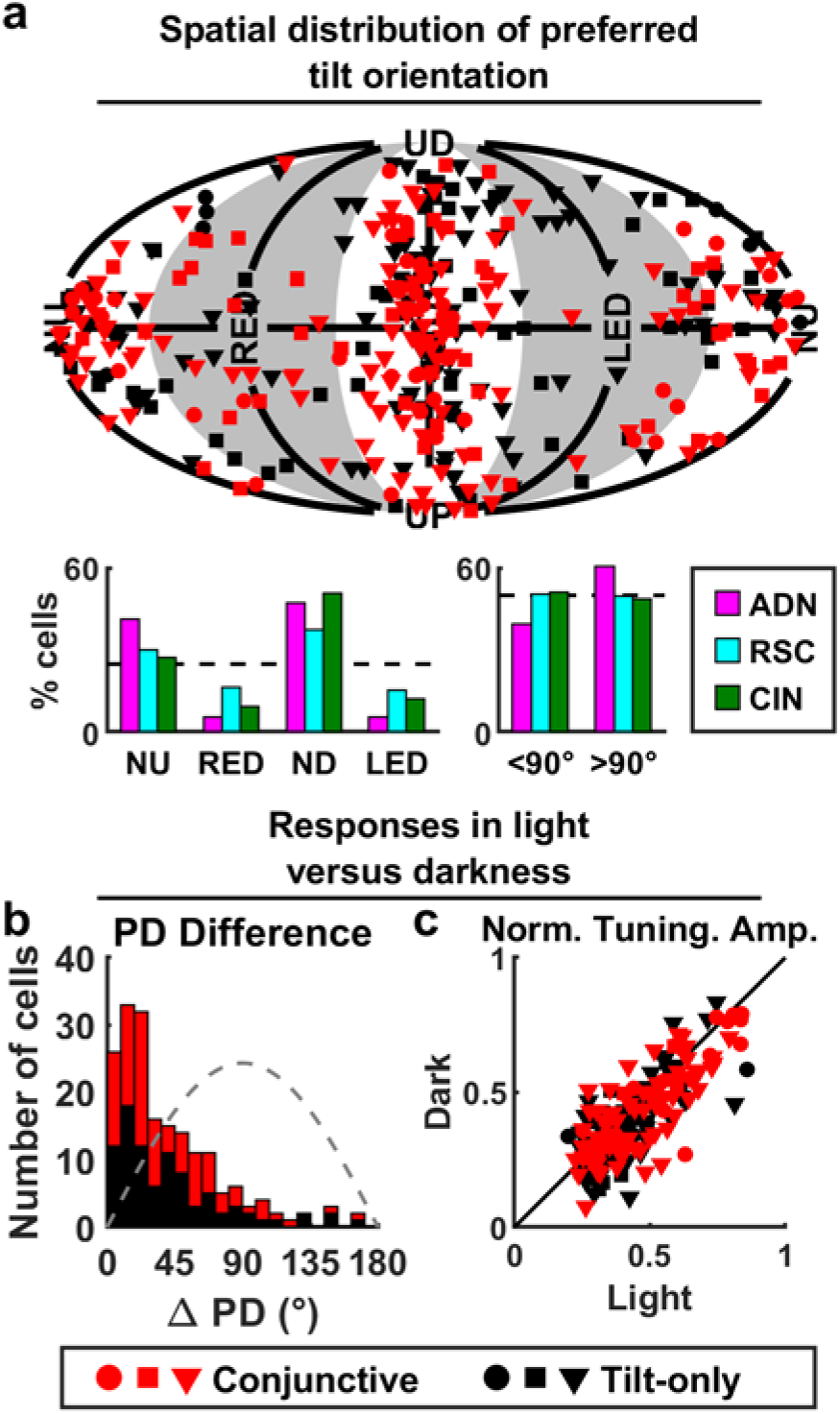
Summary of 3D tilt tuning (based on full tuning curves measured in the rotator). **(a)** Top: Distribution of tilt PDs. Red: Conjunctive (azimuth and tilt) cells; Black: tilt-only cells. Circles, squares, triangles: ADN, RSC, CIN, respectively. Bottom: Number of cells with PD in the roll (RED/LED) or pitch (ND/NU) plane; and in upper (<90° tilt) or lower (>90° tilt) hemisphere, color-coded separately for each area (AND, CIN, RSC). **(b)** Distributions of absolute difference in tilt PD for light vs. darkness for conjunctive (red) and tilt-only (black) cells. Gray dashed line: expected distribution if PDs are independently distributed on a sphere. **(c)** Comparison of tilt peak-to-trough normalized tuning amplitude computed from Gaussian fits (Methods) in darkness vs. light. Red: Conjunctive (azimuth and tilt) cells; Black: tilt-only cells.

A fundamental property of traditional HD cells is that their tuning persists in darkness (Taube, 2007; Peyrache et al., 2015). To test whether this property also characterized tilt tuning, we recorded the rotation responses of 186 (23 ADN; 30 RSC; 133 CIN) tilt-tuned cells in complete darkness. Angular differences between PDs recorded in light and darkness were close to zero (PD difference <45° in 122/186 cells, Kolmogorov-Smirnov test versus the expected distribution if PD were uniformly distributed: p = 10^−7^ in all areas, **Fig 3b**). In addition, the tilt modulation amplitude was highly correlated between light and dark conditions (**Fig. 3c**, Spearman correlation: ADN: r = 0.8, slope = 0.9; p < 10^−5^; RSC: r = 0.81, slope = 0.96; p < 10^−7^; CIN: r = 0.71, slope = 0.9; p < 10^−10^), although slightly lower in darkness in CIN (paired Wilcoxon test: p < 10^−4^; p>0.1 in other areas). These findings suggest that tilt tuning curves have similar properties as azimuth-tuning, further supporting the hypothesis that they represent different components of the same compass. Similar findings were also reported in gravity-tuned cells in the monkey anterior thalamus (Laurens et al. 2016), suggesting that these tilt-tuned neurons may be found along a broad range of animal evolution.

In further agreement with Laurens et al. (2016), a small fraction of cells also responded to tilt or azimuth velocity (**Suppl. Fig. 11**). In addition, tilt tuning in the rotator could be reproduced using traditional single-axis rotations like pitch and roll (**Suppl. Fig. 12**). Thus, tilt tuning is anchored to allocentric space, independent of the exact motion trajectory. Finally, we verified that 3D tilt tuning curves were highly reproducible across repetitions of the rotation protocol across different days (**Suppl. Fig. 13**).

### Tilt tuning is anchored to gravity

The invariance of tilt tuning in light and dark conditions (**Fig. 3b,c**), and across setups (**Fig. 2d**), supports the hypothesis that gravity could represent the allocentric reference upon which tilt tuning is based. To more explicitly test this hypothesis, we recorded 128 (22 ADN; 27 RSC; 79 CIN) tilt-tuned cells with the 3D passive re-orientation protocol after tilting the rotator and visual surround together 60° (**Fig. 1j**, magenta rotation axis). This manipulation dissociated the reference frame defined by the visual cues inside the sphere from the allocentric orientation defined by gravity (**Suppl. Fig. 14a-g**). We compared the cells’ tilt tuning curves recorded with the rotator upright (**Fig. 4a**) and when tilted (**Fig. 4b,c**). For the latter, tuning curves were computed in (i) a visual reference frame (**Fig. 4b**) and (ii) a gravity reference frame (**Fig. 4c**). If a neuron’s tuning curves measured with the system upright and tilted were identical when expressed in a gravity reference frame, then gravity served as the response anchor constant. If the tuning curves were identical when expressed in a visual reference frame, then vision was the response constant (see **Suppl. Fig. 14h**).

**Figure 4:**
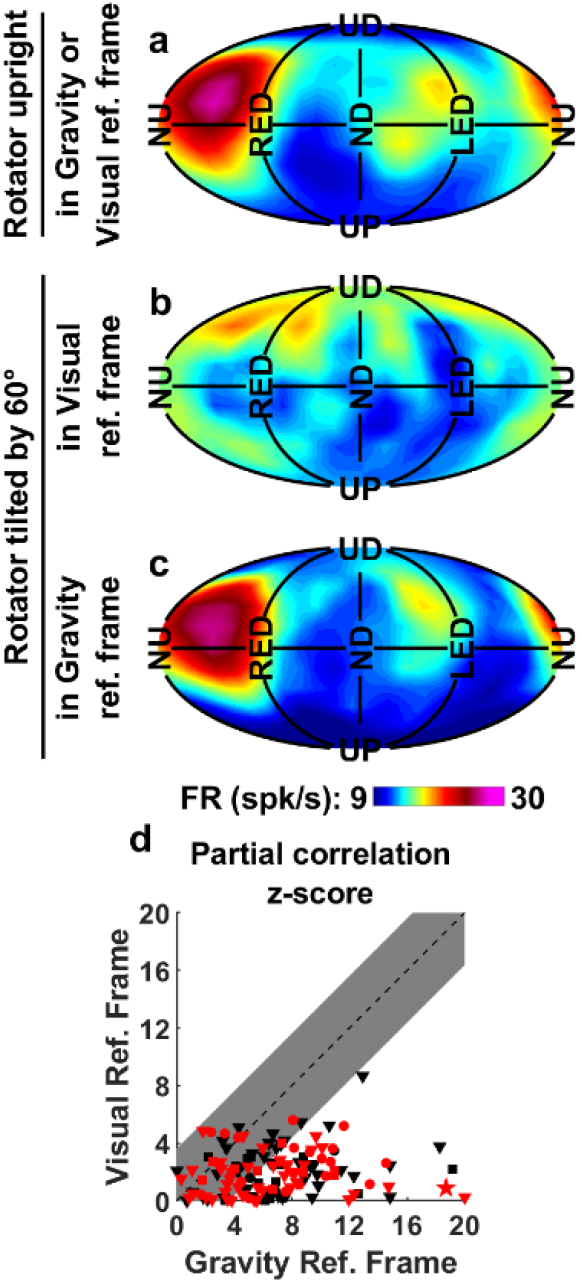
Tilt tuning is anchored to gravity. **(a-c)** Tilt tuning curves of one tilt-only example cell obtained from responses in the rotator with the spherical enclosure either **(a)** upright or **(b), (c)** tilted 60°. For the latter, tuning curves are computed in a visual *(b)* or gravity (*c*) reference frame. The visual reference frame tuning curve *(b)* is markedly distorted and attenuated relative to the one measured with the rotator upright *(a)*, unlike the tuning curves computed in a gravity reference frame *(c)*. Correlation coefficients: 0.02 (visual frame) and 0.89 (gravity frame). **(d)** Comparison between the z-scored partial correlation coefficients for Visual and Gravity reference frames across cells. Red: Conjunctive (azimuth and tilt) cells; Black: tilt-only cells. Circles, squares, triangles: ADN, RSC, CIN, respectively. Star: Example cell in (a-c). Shaded area: p>0.01.

As illustrated with the example cell in **Fig. 4a-c**, and with summary data (**Fig. 4d**), tilt tuning curves were better conserved when expressed relative to gravity cues, rather than visual cues. In fact, 76/128 cells (14/22 ADN; 15/27 RSC; 47/79 CIN) were significantly (p<0.01) more correlated to the gravity reference frame than the visual frame, and the remaining 52 cells were not significantly more correlated in either frame. Thus, although the local visual environment was identical between the upright and 60° tilt conditions, the tilt tuning followed (i.e., was invariant to) the orientation of the head relative to gravity. These findings are identical to tilt-tuned cells in the macaque anterior thalamus (Laurens et al., 2016).

### Azimuth tuning in 3D

To investigate how tilt (gravity) and azimuth components work together to encode 3D head orientation, we first questioned how to define azimuth when the head tilts away upright. The brain may simply project head direction onto the earth-horizontal (EH) plane and encode azimuth in that plane (**Fig. 5a**). Alternatively, it may represent azimuth in a compass affixed to the head-horizontal plane by tracking head rotations in this plane (yaw; **Fig. 5b**, cyan), ignoring other movements (Yaw-only model, YO, Shinder and Taube 2019). However, this possibility should be excluded on theoretical grounds, as it would not maintain allocentric invariance when the mouse locomotes in 3D (Page et al., 2017; Laurens and Angelaki, 2018). For instance, when completing the trajectory in **Fig. 5b** (red), the compass would register only 3 right-angle turns, i.e. 270°, when the head returns to its initial position. To maintain allocentric invariance when the head faces a horizontal direction (i.e. anywhere on surface 1, when facing −90/90° surface 2, or −180/180° on surface 3 in **Fig. 5b**), a head-horizontal compass must use a “dual updating rule” which includes both head-horizontal and earth (gravity)-horizontal rotations (**Fig. 5b**, green). The resulting compass was named a “tilted azimuth (TA)” frame (**Fig. 5c; Suppl. Fig. 15a**; see also Page et al. 2017; Laurens and Angelaki, 2018). Thus, in summary, we emphasize that the YO compass loses allocentric invariance during 3D motion, even when returning to upright (**Fig. 5b**). In contrast, EH and TA frames remain invariant and are identical when the head is upright (as the head-horizontal and EH planes are aligned) but differ when the head tilts (Laurens and Angelaki, 2018, **Suppl. Movie. 6**).

**Figure 5:**
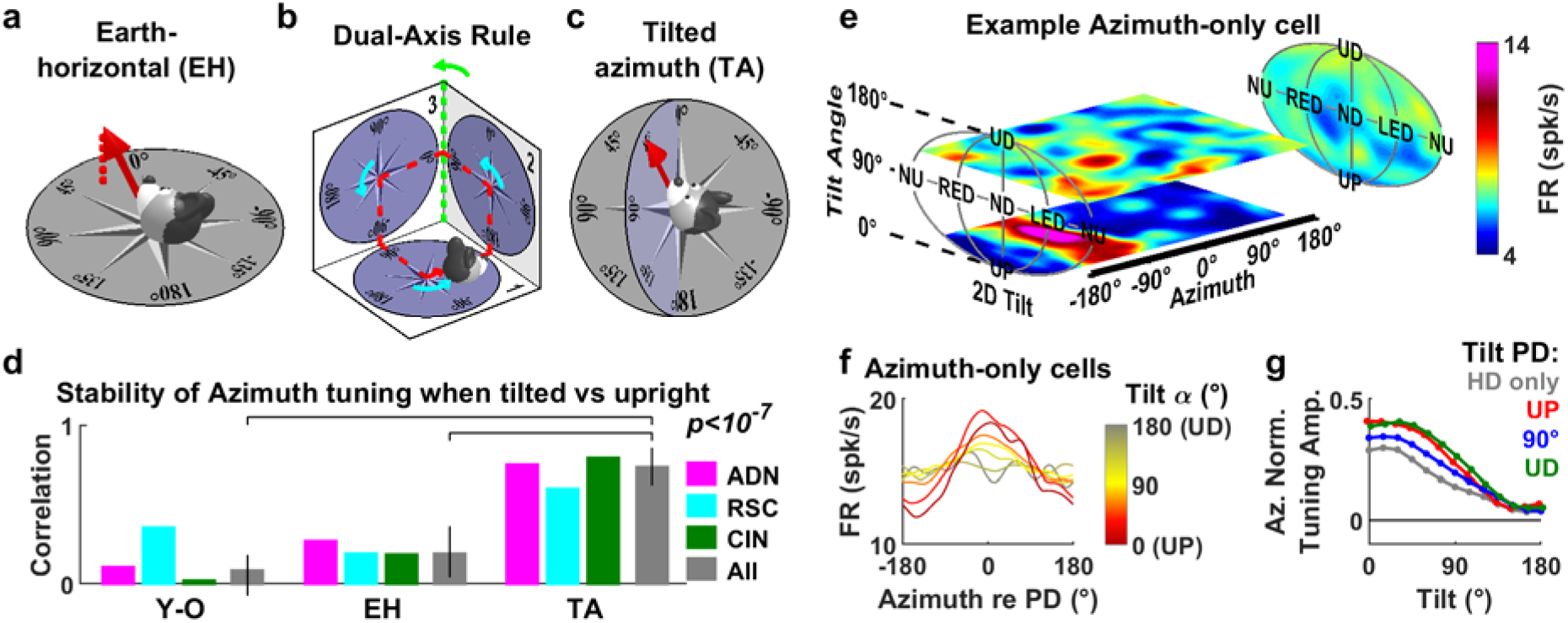
Encoding of azimuth is also influenced by gravity. **(a)** Illustration of the earth-horizontal (EH) reference frame, where azimuth direction is projected onto the earth-horizontal plane (dashed red line). **(b)** Dual-axis rule for updating an azimuth compass in a head-horizontal frame, illustrated by an example trajectory (red) where the head travels in 3D across three orthogonal surfaces (numbered 1 to 3). Head azimuth is updated when the head rotates within one surface (yaw, cyan arrows) or when it rotates in the earth-horizontal plane (green arrow when transitioning from surface 2 to 3). If azimuth is updated by yaw rotations only (yaw-only frame, YO; Shinder and Taube, 2019), the compass will not maintain allocentric consistency since it will register only 270° when the head returns to its initial position. **(c)** Tilted azimuth (TA) reference frame (Laurens and Angelaki, 2018), where head direction is measured in a compass co-planar with the head-horizontal plane but aligned with the EH compass along the line intersecting the two planes (here, the 0-180° line). The head has the same 3D orientation in (a) and (c) but its azimuth is different in the two frames (24° in EH frame and 45° in TA frame). Updating a TA compass must follow the dual-axis rule in (b). **(d)** Stability of azimuth PD in YO, EH and TA frames. For each cell, we computed the correlation between the azimuth tuning curves when the head was upright (<45° tilt) or tilted (>60°). Correlations are shown separately for each frame and for each recorded area (colored bars) or all cells pooled together (grey bars). **(e)** 3D tuning curve of an example azimuth-only cell. Horizontal slices of the tuning curves (at tilt angles of 30° and 110°) are shown (see animations in **Suppl. Movie 7** for full tuning curve). The average tilt tuning curve (across all azimuths) is shown on the right panel. **(f)** Average azimuth tuning curve of all azimuth-only cells that maintained their azimuth tuning in the rotator (n=10), averaged across all tilt orientations (γ) and computed at tilt angles (α) ranging from 0 to 180°. **(g)** Average normalized tuning amplitude of the azimuth tuning curve as a function of tilt angle, computed for azimuth only cells (n=10, grey) and conjunctive cells with preferred tilt directions near upright (<75° tilt, red, n=16), near 90° tilt (±15°, blue, n=22) and near upside-down (>105°, green, n=15).

The data collected during the 3D motion protocol in the rotator allows quantitative testing of the YO, EH and TA azimuth hypotheses. First, we expressed azimuth in all 3 frames and tested whether cells were significantly tuned when the 3D trajectory brought the head close to upright (<45° tilt). As predicted, almost none of the cells observed exhibited significant tuning in a YO frame (6/267 tuned cells, consistent with false positive at p = 0.01). In contrast, 63/267 cells were tuned as predicted when azimuth was expressed in either the EH or TA frames (ADN: n=17; RSC: n=7; CIN: n=39).

Second, when azimuth is expressed in the appropriate coordinate frame, the cell’s azimuth PD should be invariant at all head tilts (**Suppl. Fig. 15b,c,d**). To test this, we compared the cells’ azimuth tuning near upright (<45° tilt) or when tilted (>60° tilt) (**Fig. 5d**). We observed that the neurons’ tuning curves were highly correlated when expressed in a TA frame (median: 0.73; [0.63-0.86]CI), but significantly less so when expressed in a EH frame (0.19; [0.04-0.37] Cl). In addition, correlations were near zero when expressed in a YO frame (0.08, [-0.07-0.18]CI). We verified (3-way ANOVA, difference between reference frames: p<10^−10^) that correlations were similar across recorded areas (p=0.6) and tilt-tuned or non-tilt tuned cells (p=0.2).

**Figure 6:**
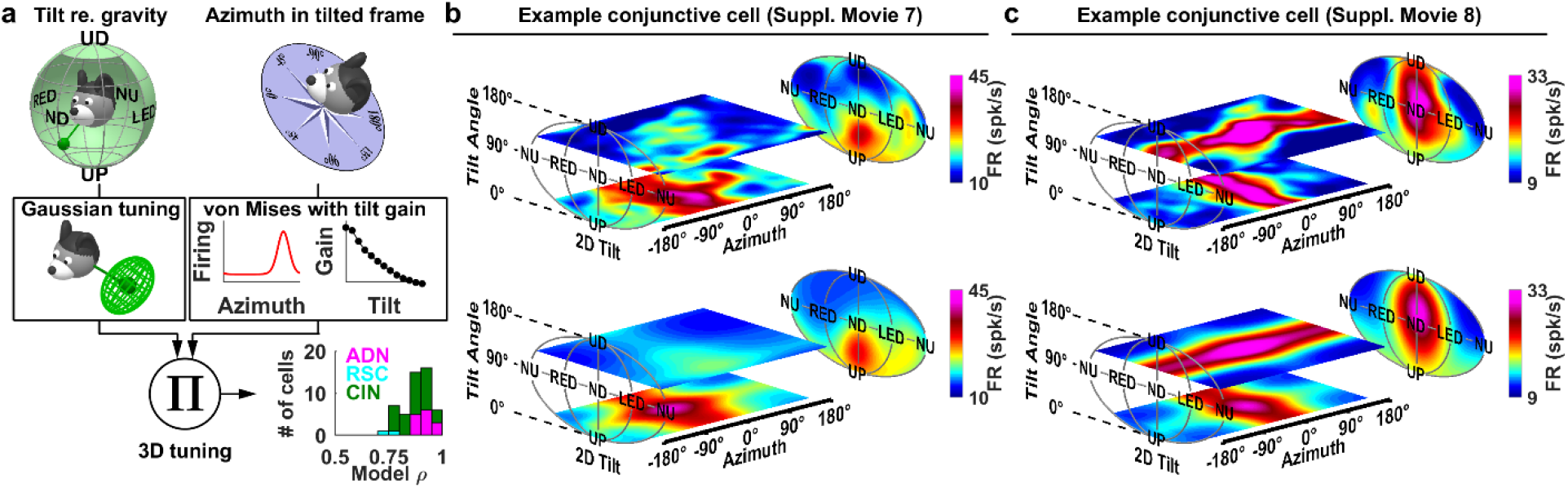
Modelling 3D responses. **(a)** Separable model overview. 3D tuning curves are the product of a 2D tilt tuning curve and a 1D azimuth tuning curve. Tilt tuning curves are produced by integrating gravity signals through Gaussian tuning functions (**Suppl. Fig. 9,10**). Azimuth tuning is expressed in a TA frame, and modelled as von Mises distributions combined with a tilt-dependent gain factor (**Fig. 5g**). Inset: Distribution of the model’s coefficient of correlation (ρ) across areas. **(b,c)** Experimentally measured 3D tuning curves (top) from two conjunctive cells that maintain their azimuth tuning in the rotator and fitted tuning curves (bottom), represented as color maps in 3D space (animated in **Suppl. Movies 8,9**).

It was also noted that, regardless of coordinate frame, the PD response amplitude of azimuth tuning often decreased when the animal was tilted beyond 90° from upright (**Suppl. Movie 6-9**). As illustrated with an example Az-only cell in **Fig. 5e** (animated curve in **Suppl. Movie 7**), azimuth tuning was strong, with a clear PD at γ = −85° for small tilt angles (lowest portion of the example cell’s tuning curve). However, response at large tilt angles (i.e, upper portion of the 3D tuning curve, **Fig. 5e**) did not exhibit any azimuth tuning. This finding was consistent for all Az-only cells: the average HD tuning curve (aligned to peak at PD=0°) had a higher modulation when computed for head tilts close to upright (**Fig. 5f**, red) and almost no modulation close to upside-down (**Fig. 5f**, grey). Thus, for Az-only tuned cells, the response amplitude of azimuth tuning was dependent on the tilt angle, even though the cells were not tuned to tilt.

That the average azimuth normalized tuning amplitude decreased monotonically with tilt angle was not only limited to Az-only cells (**Fig. 5g**, grey). Notably, the azimuth tuning amplitude of conjunctive cells decreased in a similar manner, irrespective of whether the cell’s tilt tuning favored upright orientation (**Fig. 5g**, red), intermediate tilt (i.e. 90°; **Fig. 5g**, blue) or upside-down (**Fig. 5g**, green). Normalized tuning amplitudes for all azimuth tuned cells were affected by tilt angle (two-way ANOVA, p<10^−10^) and varied between groups of cells (p<10^−4^); however there was no significant interaction effect (p=0.8), indicating that the azimuth tuning of all cells was equally affected by tilt. We conclude that HD cells encode azimuth in a TA reference frame, and that their azimuth response decreases when the head tilts away from upright, irrespective of tilt tuning.

### Tilt and azimuth tuning components of 3D compass follow multiplicative interaction

In line with experimental results and to gain further insight into the structure of 3D tuning, we created a 3D HD model that incorporates the following properties (**Fig. 6a**): (1) tilt tuning curves are generated by integrating gravity signals into Gaussian tuning functions (**Suppl. Fig. 9**), (2) azimuth-tuned cells encode TA with a tilt-dependent gain, independently from tilt tuning, (3) tilt and azimuth tuning interact multiplicatively (an additive model performed worse; Methods).

We found that these properties composed a model that could fit the tuning curves of conjunctive cells well (n=16 ADN, 4 RSC, 33 CIN; median ρ=0.88, [0.86-0.91] Cl), as illustrated with two examples (**Fig. 6b,c; Suppl. Movie 8,9**). The peak tilt response of the first example cell occurred at a tilt angle α = 42°, at which azimuth tuning (PD=-27°) had not yet attenuated. Consequently, the cell exhibited tilt and azimuth tuning simultaneously at this tilt angle, resulting in a preferred 3D orientation, visible as a region of maximum firing on both the measured (**Fig. 6b**, top) and fitted (**Fig. 6b**, bottom) curves. This type of tuning was characteristic of conjunctive cells with PDs close to upright.

The second example cell exhibited a PD at a large tilt angle (α = 105°), where azimuth tuning had already substantially decreased. As a consequence, the cell appeared azimuth-tuned at small tilt angles, where tilt tuning was minimal (**Fig. 6c**, lower horizontal plane) and tilt-tuned at large tilt angles (**Fig. 6c**, upper horizontal plane). This type of tuning curve was characteristic of conjunctive cells with a large preferred tilt angle.

Further, the same model properties also quantitatively described 3D tuning curves for azimuth-only (median ρ=0.75, [0.7-0.8] Cl) and tilt-only cells (median ρ=0.87, [0.85-0.88] Cl). Here again, model fits were significantly lower when 3D curves were computed with azimuth in an EH frame (**Suppl. Fig. 16**), confirming that the TA frame captures the cell’s response better than either the EH or YO frames could.

## Discussion

In summary, these findings demonstrate that HD cells in two areas of the mouse navigation system, as well as their output fiber bundle (CIN), are tuned in 3D. HD cells were observed to encode 2D tilt either in isolation or conjunctively with 1D azimuth. Azimuth and tilt tuning appear to represent two independent streams of information converging onto HD cells and the probabilities of individual cells being tuned to tilt and azimuth are nearly independent (**Fig. 2**). In addition, the spatial properties of azimuth tuning are independent of tilt tuning (**Fig. 5g**), and the two are separable; i.e. a cell’s entire 3D head orientation tuning curve can be computed given its tilt and azimuth tuning. Finally, azimuth tuning is anchored to visual cues (Taube 2007) and tilt tuning to gravity. At the same time, azimuth tuning is encoded in a tilted reference frame, as suggested in previous studies (Page et al. 2017; Laurens and Angelaki, 2018) which is defined by integrating 3D head rotations and gravity (Page et al. 2017, Laurens and Angelaki 2018) and not only rotations in the head horizontal plane (Calton and Taube 2005; Shinder and Taube 2019). Furthermore, the amplitude of azimuth tuning is modulated by gravity.

A recent study by Shinder and Taube (2019) concluded that HD cells encode only azimuth computed in 1D by integrating rotations in the head horizontal (yaw) plane. However, in our assessment (Laurens and Angelaki 2019), this study is in fact supportive of the tilted azimuth model, although it wasn’t directly tested in that work. Furthermore, Shinder and Taube (2019) reported no evidence of tilt tuning. However, this study is inconclusive because their interpretation of the data confounds the effects of possible tilt tuning with the fact that azimuth tuning strength is reduced as a function of tilt angle (**Suppl. Fig. 17**, see also Laurens and Angelaki 2019), a problem we circumvented by testing responses in the full 3D space. Note that Shinder and Taube observed that azimuth tuning vanishes in inverted rats, similar to (Calton and Taube 2005) and to our results. Altogether, Shinder and Taube (2019) concluded that HD cells are essentially 1D because their experimental manipulations qualitative analyses were insufficient to reveal the 3D tuning properties identified here.

The 3D model shown here to quantitatively account for the 3D orientation tuning of both azimuth-tuned and conjunctive HD cells is compatible with the toroid topology proposed for HD cells in bats (Finkelstein et al. 2015) when azimuth is expressed in a TA frame and tilt is restricted to the pitch plane (**Suppl. Fig. 18**). Tilt preferred directions are not uniformly representing 3D space, as the pitch plane is over-represented compared to the roll plane, consistent with previous findings in both macaques and bats (Finkelstein et al. 2015; Laurens et al. 2016). In fact, as summarized above (see also quantitative analysis in **Suppl. Fig. 17**), tilt tuning may also exist in rat HD cells (see also Laurens and Angelaki 2019).

We conclude that 3D tuning may be a ubiquitous feature of the mammalian HD system. We suggest that the denomination ‘head direction cell’ should also apply to tilt-tuned and conjunctive cells as well as previously described azimuth-tuned cells. Thus, the proportion of HD cells is much higher than previously reported, based upon response conditions that only considered the horizontal plane.

Our study is the first to record HD cells directly in the cingulum fiber bundle that conveys ADN and RSC projections to parahippocampal regions (ADN and RSC projections), RSC (ADN projections) and cingulate cortex (RSC projections) (Domesick 1970; van Groen and Vyss 1990,1995; Bubb et al. 2018). Although recordings of axonal spikes with tetrodes are uncommon, they have been performed previously (Robbins et al. 2013). Furthermore, histology clearly demonstrates that recordings occurred in white matter (**Suppl. Fig. 3c**) and units recorded in the CIN exhibited short spike duration consistent with axonal spikes (**Suppl. Fig. 19b**; Barry 2015). Furthermore, as the CIN contains projections from the ADN and RSC, the properties of tilt and azimuth responses recorded there were consistent with those in ADN and RSC. The existence of tilt signals in the CIN supports the notion that these signals are communicated between various regions of the limbic system and therefore form an inherent component of the spatial navigation system.

A recent imaging study (Kim and Maguire 2018) indicates that the human RSC encodes pitch orientation in a virtual navigation task, although the ADN was found to encode mainly azimuth. It is possible that visually-driven tilt signals arise in the RSC in a virtual environment where visual, but not inertial gravity cues, are present. RSC is involved in visual processing, possibly in transforming visual landmark form an egocentric to an allocentric reference frame (see Clark et al. 2018, Mitchell et al. 2018 for reviews). This raises the possibility that the RSC may use gravity-referenced tilt signals to transform visual signals in 3D.

We have demonstrated that tilt signals are anchored to gravity. Indeed, gravity is a veridical vertical allocentric cue, that largely dominates vision in human verticality perception (Vingerhoets et al. 2009). Further, gravity sensing represents one of the most ubiquitous sensory modalities of terrestrial living organisms (Sack 1991, Bender and Frye 2009). The origins of the gravity signal are likely multisensory, involving proprioceptive and vestibular inputs (Clemens et al, 2011; Foisy and Kapoula 2018), as well as computations in central vestibular areas (Laurens and Angelaki 2011; Laurens et al. 2013a,b).

Prominent views of azimuth-tuned HD cells posit that they form a neuronal attractor that can memorize azimuth in the absence of sensory inputs and is updated by integrating rotation signals (Clark and Taube 2012, Peyrache et al. 2015, Laurens and Angelaki 2018), although some HD cells in the RSC (Jacob et al. 2017), and parahippocampal regions (Kornienko et al. 2018) may not contribute to the attractor network. Establishing whether tilt-tuned cells also form an attractor will be challenging, especially since gravitational input that anchors tilt tuning is not easily altered, unlike the visual input that anchors azimuth tuning. Alternatively, there may not be a gravity attractor, but rather gravity signals may be computed directly by central vestibular pathways, where the mathematical challenge of integrating 3D inertial cues (Green et al. 2005) is already solved (Laurens et al. 2013b), thus providing a computationally simpler solution for a 3D orientation compass. Exploring neuronal computations that underlie 3D orientation coding in the limbic system and their relationship with pre-limbic computations in the brainstem, cerebellum, thalamus and hypothalamus must be targeted in future studies.

Tilt and 3D orientation tuning had previously only been identified in aerial (bats) and tree-dwelling (macaques) species, raising the question of whether a 3D compass would be ethologically relevant to rodents. Although laboratory mice (*Mus musculus*) and rats (*Rattus norvegicus*) are primarily land-dwelling, they exhibit a rich 3D behavioral repertoire (Makowska and Weary 2016; Mimica et al. 2018) in the wild living in extensive underground burrows (Schmidt and Fischer 2011) and easily learn 3D spatial orientation tasks (Wilson et al. 2015). Furthermore, laboratory mice and rats are physiologically related to tree-dwelling rodents (Arregoitia et al. 2017), including other muroids (e.g. harvest mice, *Micromys minutus)* and non-muroids (e.g. squirrels). It is therefore not surprising that rodents, like bats (Finkelstein et al. 2015) and likely macaques (Laurens et al. 2016) and humans (Kim and Maguire 2018), possess a three-dimensional compass, whose properties may be phyla-independent.

## Authors contributions

JL designed and supervised experiments, analyzed data and wrote manuscript. JN, AMA, HC performed experiments and analyzed data. JDD supervised experiments. DEA designed and supervised experiments and wrote manuscript.

## On-line Methods

### Animals

A total of 10 male adult mice (C57BL/6J), 3-6 months old, were used in this study (**Suppl. Table 1**). Animals were prepared for chronic recordings by implanting a head-restraint bar and a microdrive/tetrode assembly under general anesthesia (Isoflurane) and stereotaxic guidance. Two skull screws were implanted in the vicinity of the target region, and a circular craniotomy (~1.5mm diameter) was performed above the target region. Animals were single-housed on a reversed [12/12] light/dark cycle. Experimental procedures were conducted in accordance with US National Institutes of Health guidelines and approved by the Animal Studies and Use Committee at Baylor College of Medicine.

### Neuronal recordings

Neurons were recorded using 6 (mice AA1/AA2) or 4 (all other mice) tetrode bundles constructed with platinium-iridium wires (17 micrometers diameter, polyimide-insulated, California Fine Wire Co, USA) and platinum-plated for a target impedance of 200kΩ using a Nano-Z (Neuralynx, Inc) electrode plater. Tetrodes were cemented to a guide tube (26-gauge stainless steel) and connected to a linear EIB (Neuralynx EIB/36/PTB). The tetrode and guide tube were attached to the shuttle of a screw microdrive (Axona Ltd, St Albans, UK) allowing a travel length of ~5mm into the brain.

The stereotaxic coordinates for each tetrode implant was based upon Bregma as a reference point. The coordinates used to target both the ADN and the CIN were 0.2 mm posterior and 0.7 mm lateral to Bregma. The granular/dysgranular RSC were targeted by implanting 2.0 mm posterior and 0.07/0.7 mm lateral to Bregma respectively.

#### Postmortem verification of electrode sites

At the end of the study, brains were removed for histological verification of electrode location. The animals underwent transcardial perfusion with 4% paraformaldehyde (PFA). The brains were postfixed in 4% PFA and then transferred to 30% sucrose overnight. Brain sections (40μm) were stained (Nissl or neutral red staining), and examined using bright-field microscopy to localize tetrode tracks (**Suppl. Figure 2**). Photographs of histological slides were corrected for brightness, contrast, gamma and color balance.

#### Firing properties

We confirmed that the distribution of average firing rates and CV2 were similar for all cells in the CIN, ADN, and RSC (**Suppl Fig. 19a**). Furthermore, we observed that most spikes recorded in CIN had small (<0.3ms) trough to peak durations, indicative of axonal spikes (**Suppl Fig. 19b**).

### Experimental apparatus

#### Freely moving recordings

In order to identify traditional HD cells, we first recorded as mice explored freely in a circular arena (50 cm diameter, 30 cm height; **Fig. 1b**). The walls of the arena were white with a 45° black card to provide a visual orientation cue. To record tilt tuning in freely moving mice, the arena was replaced by a movable platform that was constructed by mounting an oblong nylon mesh (20×30cm, 1.5cm mesh) onto a manually operated three-axis gimbal system; **Suppl. Fig. 4a**. The system was placed at the center of a large cylinder (130 cm diameter, 2 m height), its door was left open during recording to provide a visual landmark and to allow the experimenter to monitor each mouse. In both systems, neuronal data were acquired at 22 kHz using a MAP system (Plexon Inc.). The microdrive’s EIB was plugged to a tethered head stage that included two LEDs (one red and one infra-red, 4 cm apart) for optical tracking (Cineplex, Plexon Inc.). In addition, mice’s head were equipped with a digital 6-degree-of-feedom inertial measurement unit (IMU; SparkFun SEN-10121) for measuring head tilt relative to gravity. Perspective effects that could affect optical tracking when the head tilted away from horizontal were corrected based on the IMU data.

#### Motorized rotator

To measure the 3D orientation tuning using a uniform representation of tilt angles, we tested animals using a motorized rotator. It also allowed us to separate visual from gravity representations. We gently restrained each mouse’s body and fixed its head rigidly, and placed it in the center of a rotation simulator (**Suppl. Fig. 6a**) composed of a motorized three-axis motion system (Axes I-III in **Suppl. Fig. 6a**) inside a visual surround sphere (1.8 m diameter) (Acutronics Inc., Switzerland). The inside of the sphere was painted in white, with three horizontal lines of dots (10° diameter, 30° spacing) to provide horizon and optokinetic cues. Three vertical LED stripes were placed 22.5° apart to provide a horizontal orientation cue. A fourth rotation axis (Axis IV) allowed tilting the rotator and the sphere together (sphere door closed). Neuronal data were acquired at 30 kHz using a neural data acquisition system (SpikeGadget, San Francisco, California). The position of the rotator’s axes (and therefore the 3D orientation of the head) was measured with potentiometers installed in each rotation axis and digitized at 833 Hz.

All recorded data was organized in a custom-made database using Datajoint (Yatskeno et al. 2015)

### Experimental protocols (see Suppl. Table 2)

#### Experiment 1: Characterization of HD tuning in the arena

We recorded neuronal responses during five 8-minute sessions. A first recording session was performed in light (*Experiment 1-L0*). We then performed the other protocols described below on a moving platform and rotator, before returning the mouse to the same arena and performing three separate 8-min sessions, first in light (*Experiment 1-L1*), then in darkness (*Experiment 1-D*), then we repeated a session in light (*Experiment 1-L2*).

#### Experiment 2: Tilt tuning in freely moving animals

We recorded neural responses when mice walked freely on a platform. Recordings were performed in 5-min blocks during which the setup’s axis II and III were fixed. Within a single block with the platform tilted, freely movements on the platform’s surface changed the mice’s head azimuth (Az) and tilt orientation (angle y) together, and these variables are therefore correlated (**Suppl. Fig. 4b**, yellow, magenta). Rotating the base (rotation along the blue arrow) between blocks added an offset to azimuth, while leaving the range of head tilt unchanged (e.g. **Suppl. Fig. 4b**, yellow vs. magenta). This manipulation allowed coverage of all possible head azimuth and tilt orientations (plane in **Suppl. Fig. 4b**), thereby allowing coverage of a large portion of 3D space relatively uniformly (up to α=60°; **Suppl. Fig. 4c**).

We perform one additional manipulation of the space covered by the animal: Half-way through each block, the platform was rotated using axis I (**Suppl. Fig. 4a**). This manipulation served as control for the following potential confounding factor: As long as only Axis III is operated, then local azimuth on the platform is anchored to gravity, e.g. the same side of the platform is always placed downward. Therefore, if a cell’s firing was anchored to the azimuth on the platform itself, and not to the tilt, its response could be misinterpreted as a tilt response. Changing Axis I multiple times within each block randomizes local azimuth relative to tilt, which prevents this potential confound.

We performed 17 blocks (~68 min) with the following organization: (1) one block where the platform was horizontal (duration: 8 min), (2) eight blocks where the platform was tilted 45° and the base was rotated in steps of 45° (duration: 2.5 min) and (3) eight blocks where the platform was tilted 65° and the base was rotated in steps of 45° (duration: 5 min). Note that mice tended to upright their head, therefore tilting the mesh 45° and 70° resulted in average head tilts of ~35° and 60°, respectively (e.g., **Suppl. Fig. 4c**). Together, these 13 blocks allow a relatively uniform sampling of 3D head orientation (at tilt angles up to ~60°) while mice were unrestrained and locomoting freely.

To ensure that tilt space was adequately sampled, we computed the occupancy distribution d (i.e. the time spent) across 73 tilt positions (uniformly distributed in tilt space for up to 60°). Next, we computed the entropy of d E(d)=-Σp(d).log_2_(p(d)), ranging from log_2_(73)=6.19 (uniform distribution) to 0 (if the mouse occupies a single point). We excluded cells where E(d)<5.6, which corresponds to mice sampling less than 2/3 of the tilt space. 45% of recorded cells (not counted in **Suppl. Table 1**) were excluded based on this criterion.

#### Experiment 3: Three-dimensional tuning in the rotator

The rotator was programmed to scan 3D rotation space uniformly using preprogrammed trajectories that sample 200 head tilt orientations uniformly (**Suppl. Fig. 6b**, red; **Suppl. Movie 3**); the distance between adjacent points being ~15°. We computed four distinct trajectories (no overlap, **Suppl. Fig. 6c**, different colors), each of which visited all points once, and in different order. Trajectories travelled through each point in a straight line at a constant velocity (30%) and changed direction between points (**Suppl. Fig. 6b,c**). All trajectories were replayed forward and backward. This technique ensures that the 2D space of head tilt is covered uniformly. While the desired head tilt is achieved by controlling the two innermost axes (I and II), azimuth is varied by rotating axis III (outer) of the rotator at a constant velocity (±15°/s; **Suppl. Fig. 6d**, red; the velocity is reversed every 4 rotations). During the trajectory, mice always faced at least 90° away from the second axis (black in **Suppl. Fig. 6a**) to ensure that the visual field in front of the mouse is not obstructed. We performed the following variants of the protocol: (i) with the LED stripes (placed inside the visual enclosure) on (*Experiment 3-L*), (ii) off (*Experiment 3-D*), and (iii) LED on, after the rotator and the visual enclosure were tilted en bloc 60° relative to vertical by operating Axis IV (**Suppl. Fig. 14**) (*Experiment 3-T*).

#### Experiment 4: Yaw/pitch/roll rotations

The rotator was programmed to rotate each mouse back and forth in yaw, pitch or roll at a constant velocity of 30°/s. Starting from a velocity of 0°, each movement included an acceleration phase of Is to 30°/s, then 380° of rotation at constant velocity and finally a deceleration period of 1s. To exclude any potential response to accelerations or decelerations, only data recorded during the central 360° of constant-velocity rotation period was used in the analysis.

### Data analysis

#### Neuron selection and classification

All well isolated neurons recorded during at least one foraging session in light in the arena (Experiment 1-L0, L1 or L2), and during Experiment 2 or Experiment 3-L have been included for analyses. We first classified neurons as azimuth-tuned or non-azimuth-tuned based on their responses in the freely moving arena. Neurons could also be classified as azimuth-tuned or azimuth-untuned based on their responses in the platform and rotator. However, because azimuth responses have lower amplitude in the rotator, they often didn’t reach significance level. Therefore, throughout the study, “azimuth-tuned” refers by default to the classification based on freely moving data in the arena (Experiment 1).

Similarly, neurons were classified twice as tilt-tuned or not, based on recordings on the orientable platform and in the rotator independently. As with azimuth tuning, neurons that exhibited significant tilt tuning when moving freely may not be significantly tuned in the rotator, because responses in the rotator had lower amplitude. On the contrary, some neurons that exhibited significant tilt tuning in the rotator weren’t significantly tuned when moving freely because this protocol sampled a limited range (~1/3) of head tilt. Nevertheless, differences were small, and the majority of neurons significantly tilt-tuned in one setup were also tuned to the other (**Fig. 2**).

Importantly, we confirmed that azimuth and tilt tuning in the rotator and freely moving were correlated in terms of amplitude and consistent in terms of spatial characteristics for neurons that were significantly tuned to azimuth or tilt in both experiments (**Fig. 2c,d; Suppl. Fig. 8**).

#### Tuning curves

For each recorded neuron, we computed the following tuning curves:

1. To evaluate azimuth tuning, we computed 1D azimuth tuning curves in all conditions of *Experiment 1*, in *Experiment 2* when the platform is horizontal, in *Experiment 4-Ygw*; and in data points where head tilt was less than 45° during *Experiment 3-L*.
2. To evaluate tilt tuning for *Experiment 2* and *Experiment 3-L,D,T*, data were averaged across azimuth. We also computed pitch/roll tuning curves based on *Experiment 4*.

Note that tilt tuning is different from a recent finding that RSC HD cells encode azimuth in a tilt-dependent manner (Page et al. 2017), which has been explained by a framework called ‘the dual-axis rule’ (Page et al. 2017) or ‘tilted azimuth’ (Laurens and Angelaki 2018); see **Suppl. Fig. 15**. Although tilt influences the azimuth reference frame, tilted azimuth signals do not carry any information about head tilt. Reciprocally, the tilt tuning identified here doesn’t carry any information about azimuth. Thus, 2D tilt tuning and 1D tilted azimuth encode different dimensions of 3D head orientation (see also **Suppl. Movie 1**).

#### Coordinates used to encode 3D head orientation

Head tilt was expressed in spherical coordinates (α,γ; **Suppl. Fig. 5**), where α is the tilt angle: α=0° in upright orientation (UP) and α=180° in upside-down orientation (UD); and γ encodes tilt orientation: γ = 0° and γ = 180° correspond to nose-down (ND) and nose-up (NU) tilt (pitch); γ = 90° and γ = −90° correspond to left-ear-down (LED) and right-ear-down (RED) tilt (roll). For fitting purposes (e.g. **Suppl. Fig. 9**), head tilt was also expressed in Cartesian coordinates (G_X_, G_Y_, G_Z_) that represent the orientation of the gravity vector (normalized to a length of 1) in head coordinates. Spherical coordinates are transformed into Cartesian coordinates and vice-versa by G_X_=sin(α).cos(γ); G_γ_= sin(α).sin(γ); G_z_= -cos(α); and α=acos(-G_z_); γ=atan2(G_Y_,G_X_).

Neuronal responses were evaluated by computing tuning curves, which were smoothed using Gaussian kernels with standard deviation of 15° on both azimuth and tilt. We computed 3D azimuth tuning curves in 3 different ways, by expressing azimuth in a yaw-only (Y-0), an earth-horizontal (EH) or a tilted (TA) frame, and found that the latter accounted for neuronal responses better (**Fig. 5; Suppl. Fig. 15**). Earth-horizontal azimuth is computed by defining a “forward” pointing vector N, aligned with the head’s naso-occipital axis, and encoding its orientation in an earth-fixed reference frame (i,j,k), i.e. N=(N_i_, N_j_, N_k_). Earth-horizontal azimuth is defined as the orientation of N on the earth-horizontal (l,j) plane, i.e. EHAz = atan2(N_j_, N_i_). EH azimuth can be transformed into tilted azimuth by the following equation: TA = EHAz – γ – atan2(-sin(γ),cos(α).cos(γ)).

#### Tuning curve fitting

To quantify tuning curves, von Mises and/or Gaussian functions were fitted and standard shuffling analysis was used to evaluate the statistical significance of azimuth and tilt tuning.

2D tilt tuning curves were fitted with Gaussian distributions (see **Suppl. Fig. 9,10**), where tilt was expressed in Cartesian coordinates and FR_Ti_(α,γ) = FR_0_ + A.N_M,C_(G_X_, G_Y_, G_Z_) where N_M,C_(G_X_, G_Y_, G_Z_) is a 3D Gaussian distribution centered on M and with covariance matrix C.

Azimuth tuning curves were fitted with circular normal (von Mises) distributions. Preliminary analysis revealed that the PD of azimuth tuning is maintained when the head tilts (when azimuth is expressed in a tilted frame) but that its gain changes. To account for this, we defined a tilt-dependent gain G(α) and expressed azimuth tuning as:

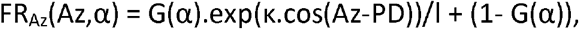

where k is the parameter of the van Mises distribution. For convenience, we normalized FR_Az_(Az,α) such that its average value across all azimuths is 1 (by setting I to the average value of exp(k.cos(Az-PD))).

Finally, we evaluated the interaction between azimuth and tilt tuning by fitted 3D tuning curves as the product of the azimuth and tilt tuning cures defined above, i.e.

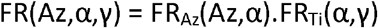

Tilt tuning curves had 11 free parameters: FR_0_, A, M (3-dimensional) and C (a covariance matrix, i.e. 6-dimensional). The normalized azimuth tuning curves had 2 free parameters (k and PD).

The tilt-dependent gain G(α) was fitted independently at 13 tilt angles α ranging from 0 to 180° by increments of 15°, resulting in 13 additional free parameters. The 3D tuning curves were computed from experimental data at 184 uniformly distributed tilt orientations and 24 azimuth orientations, i.e. 4416 points, and fitted to the 3D curve model by gradient ascent (Matlab function Isqnonlin). Note that since the average value (across all azimuths) of FR_Az_(Az,α) was 1, the average tilt tuning curve (across all azimuth) was FR_Ti_(α,γ).

We tested an additional model that assumes that azimuth and tilt tuning interact additively, i.e. FR(Az,α,γ) = FR_Az_(Az,α)+FR_Ti_(α,γ). We found that this model didn’t fit 3D tuning curves as well as the multiplicative model: its correlation coefficient was significantly lower in 33/53 conjunctive cells, and better only in 1/53 conjunctive cell (Fisher r to z transform, at p<0.01), and significantly lower at the population level (median ρ=0.85, [0.80-0.87] Cl versus 0.88, [0.860.91] Cl, p<10^−8^, paired Wilcoxon test). Therefore, we used the multiplicative model to model 3D responses in this study.

#### Quantification of tuning strength

The length of the mean vector (i.e. the normalized Rayleigh vector), ranging from 0 to 1, is used commonly to assess how strongly a cell is tuned (|R| =0, untuned cell; |R|=1, maximally tuned cell) independently from its average firing rate. It allows comparing cells with a large range of peak firing rates. Thus, azimuth tuning was quantified consistently with previous studies by computing the mean vector R = c.Σ FR(Az)*exp(-i*Az)/ Σ FR(Az), where FR(Az) was sampled at 100 positions separated by 3.6° and c = 3.6*π/180/2/sin(1.8) (Zar, 1998).

Mean vectors can be generalized to a 2D distribution by expressing tilt in Cartesian coordinates G=(G_X_, G_Y_, G_Z_) and computing R = Σ FR(G)*G/ Σ FR(G). The resulting 2D vector has a length of 1 if all spikes occur at the same tilt and 0 if spikes are distributed uniformly or symmetrically.

However, because mean vectors computed in 1D and 2D can’t be compared directly, we developed an alternative measure called normalized tuning amplitude (NTA; **Suppl. Fig. 7**). The normalized tuning amplitude of an 1D azimuth or 2D tilt tuning curve was defined based on the maximum (FR_max_) and minimum (FR_0_) firing rate, as A_N_ = (FR_max_-FR_0_)/ FR_max_. Thus, normalized tuning amplitude ranged from 1 (when a tuning curve ranged from 0 spk/s to a peak value) to 0 (when a cell was unmodulated). Note that normalized tuning amplitude measures the cell’s modulation amplitude, but not the sharpness of the tuning curve.

#### Statistical procedures to determine significant tuning

We used a shuffling procedure (Langston et al. 2010; Yartsev et al; 2011) to assess the statistical significance of azimuth or tilt tuning. Each sample was generated by (1) shifting the entire spike train circularly by a random value of at least ±10s, (2) recomputing the tuning curve, (3) performing the Gaussian fit and (4) computing the azimuth and/or tilt normalized tuning amplitude. We computed the mean value *m* and standard deviation *σ* of the normalized tuning amplitude across 100 shuffled samples. The statistical p-value of the normalized tuning amplitude NTA measured in the un-shuffled data was computed as 1-F(NTA, m, σ), where F is the cumulative Gaussian distribution with average m and standard deviation σ.

We considered azimuth or tilt tuning to be significant if (1) the p-value computed as described above was less than 0.01 and (2) the normalized tuning amplitude NTA was equal to or higher than 0.25. The second criterion was equivalent to selecting cells where the modulation was at least one third of the baseline firing rate, and was used to eliminate cells with very small but significant modulation (how this threshold compares to criteria used in other studies is analyzed and discussed in **Suppl. Fig. 7**).

We combined data from multiple repetitions of Experient 1-L to assess if cells were significantly tuned to azimuth when the animal was moving freely. We used two techniques to combine multiple repetitions: (1) we analyzed each repetition independently and computed the median p-value and amplitude across repetitions, and (2) we computed a p-value and amplitude based on data pooled across all repetitions. We tested if the values obtained with technique (1) or with technique (2) passed the criteria described above, and classified cells as azimuth tuned of they passed any of these tests. This two-technique approach was used because pooling data yields a greater statistical power but fails if the cell’s PD shifted between sessions, whereas the second approach isn’t affected by shifts in the PD. In total, 37% cells passed both tests; 6.5% passed the first test only, 7.5% passed the second test only, and 51% cells passed one or the other and were classified as azimuth-tuned.

We used data from Experiment 3-L to assess if cells are significantly tuned to tilt, by testing if the normalized tuning amplitude of the average tilt tuning curve (across all azimuth) passed the criteria described above. We also tested if cells are significantly tuned to azimuth in the rotator by computing the azimuth tuning curve based on data for up to 45° tilt and testing if its normalized tuning amplitude passes the criteria described above. In some cells, Experiment 2 or Experiment 3-L were repeated multiple times. We found that the preferred direction of tilt and azimuth tuning were stable across repetitions, and pooled data across all repetitions.

#### Tilt and azimuth velocity analysis

We performed another Gaussian curve fitting to test whether neurons carry a mixture of tilt and tilt derivative information. We expressed head tilt measured during Experiment 3-L in Cartesian coordinates (G_X_, G_Y_, G_Z_) and then computed the time derivative of the gravity vector (dG_X_/dt, dG_Y_/dt, dG_Z_/dt) as a measure of tilt velocity. Next, we fitted neuronal firing rate with FR = FR_0_ + A.N_M,C_(G_X_, G_Y_, G_Z_)+ A’.N_M’,C’_(dG_X_/dt, dG_Y_/dt, dG_Z_/dt). We computed the normalized tuning amplitude of tilt and tilt velocity and used the same shuffling method and criterion (p<0.01, A_N_>0.25) as in other Gaussian fits to assess whether cells were significantly tuned to the gravity derivative. Note that the tilt tuning curves obtained with this method were identical to those obtained in previous Gaussian fit.

We investigated whether cells encode azimuth velocity (dAz/dt) by computing azimuth velocity tuning curves for each cell (i.e. average firing rate as a function of dAz/dt) using all data from Experiment 1-L. The tuning curves were evaluated at all velocities ranging from −200 to 200°/s by increment of 20°/s and smoothed using a 8°/s Gaussian kernel. We computed the amplitude and normalized tuning amplitude of these curves, and used the same shuffling method and criterion (p<0.01, A_N_>0.25) to assess whether cells were significantly tuned.

#### Experiment 3-T

To analyze the results of Experiment 3-T, we computed neuronal tuning curves in both gravity and visual reference frames. Tuning curves in a gravity frame (curve G) were computed based on the actual tilt of the head relative to gravity. Tuning curves in a visual frame (curve V) were computed as if Axis IV of the system had not been tilted. Next, we performed a linear regression of the tuning curve recorded with the rotator upright (Experiment 3-L) and the curves G and V, computed the partial coefficients of correlation of curves G and V, and z-transformed these coefficients (with 43 d.o.f. since curves were sampled at 46 points). Statistical significance was assessed at p<0.01 if the absolute difference between the z-scored partial correlations was >sqrt(2)*2.56.

#### Experiment 4P/R

Using the full 3D tuning curve data from Experiment 3-L, we “predicted” the responses to pitch and roll rotations by sampling the 3D tuning curve at the head orientations visited during pitch and roll rotations. The pitch and roll tuning curves measured during Experiment 4 and predicted based on Experiment 3 were then fitted with 1D Gaussians, and the resulting modulation amplitudes and preferred direction were compared. Data during Experiment 3 and 4 in light and darkness were averaged.

**Supplemental Table 1:** Number of recorded cells and categories. For each mouse (and each area), the table indicates the number of cells recorded during Experiment 1 (numbers in top row), 2, 3 and 4), along with their categorization. The last column shows the total number of neurons tested for each experimental protocol.

| Animal name | H71M | H72M | I10M3 | I29M | H65M | H68M | H69M | H74M | AA1 | AA2 | Total |
| --- | --- | --- | --- | --- | --- | --- | --- | --- | --- | --- | --- |
| Region | ADN | ADN | ADN | ADN | CIN | CIN | CIN | CIN | RSC | RSC |  |
| Total | 35 | 13 | 6 | 15 | 72 | 137 | 61 | 27 | 53 | 124 | 543 |
| *Azimuth tuned* | 33 | 9 | 6 | 4 | 34 | 68 | 28 | 17 | 26 | 58 | 283 |
| *not Azimuth tuned* | 2 | 4 | 0 | 11 | 38 | 69 | 33 | 10 | 27 | 66 | 260 |
| Experiment 2 Platform | 19 | 0 | 5 | 5 | 0 | 19 | 14 | 9 | 28 | 40 | 139 |
| *Conjunctive (Az&Tilt)* | 16 | 0 | 5 | 3 | 0 | 5 | 4 | 3 | 5 | 14 | 55 |
| *Tilt-only* | 0 | 0 | 0 | 1 | 0 | 6 | 4 | 4 | 6 | 16 | 37 |
| *Azimuth-only* | 1 | 0 | 0 | 0 | 0 | 1 | 2 | 1 | 9 | 5 | 19 |
| *Not HD* | 2 | 0 | 0 | 1 | 0 | 7 | 4 | 1 | 8 | 5 | 28 |
| Experiment 3 L,D,T Rotator - in Light | 30 | 13 | 3 | 14 | 72 | 137 | 59 | 19 | 53 | 114 | 514 |
| *Conjunctive (Az&Tilt)* | 30 | 8 | 2 | 3 | 28 | 47 | 24 | 11 | 14 | 37 | 204 |
| *Tilt-only* | 0 | 3 | 0 | 7 | 23 | 45 | 20 | 3 | 15 | 43 | 159 |
| *Azimuth-only* | 0 | 1 | 1 | 0 | 6 | 21 | 3 | 2 | 12 | 17 | 63 |
| *Not HD* | 0 | 1 | 0 | 4 | 15 | 24 | 12 | 3 | 12 | 17 | 88 |
| Rotator - in Darkness | 12 (12) | 5 (4) | 2 (1) | 8 (6) | 47 (33) | 98 (68) | 43 (32) | 0 (0) | 0 (0) | 44 (30) | 259 (186) |
| Rotator - Tilted | 12 (12) | 5 (3) | 1 (1) | 8 (6) | 41 (26) | 55 (38) | 22 (16) | 0 (0) | 0 (0) | 38 (27) | 182 (129) |
| Experiment 4: yaw, pitch, roll | 0 (0) | 0 (0) | 0 (0) | 0 (0) | 40 (28) | 18 (14) | 11 (8) | 0 (0) | 0 (0) | 0 (0) | 69 (50) |

**Supplemental Table 2:**
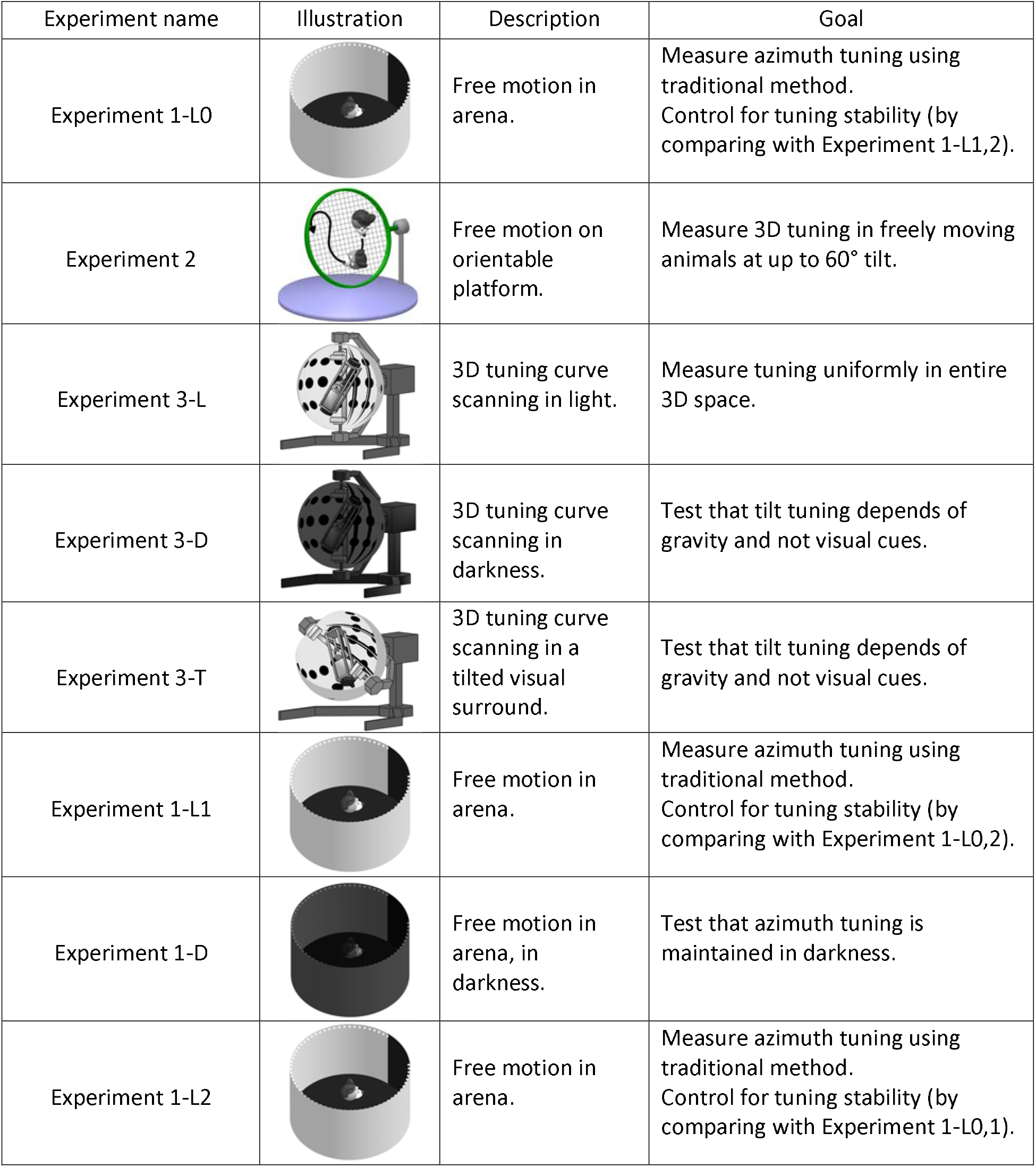
Description and order of experimental protocols. For each mouse (and each area), the table indicates

| Experiment name | Illustration | Description | Goal |
| --- | --- | --- | --- |
| Experiment 1-L0 |  | Free motion in arena. | Measure azimuth tuning using traditional method.<br>Control for tuning stability (by comparing with Experiment 1-L1,2). |
| Experiment 2 |  | Free motion on orientable platform. | Measure 3D tuning in freely moving animals at up to 60° tilt. |
| Experiment 3-L |  | 3D tuning curve scanning in light. | Measure tuning uniformly in entire 3D space. |
| Experiment 3-D |  | 3D tuning curve scanning in darkness. | Test that tilt tuning depends of gravity and not visual cues. |
| Experiment 3-T |  | 3D tuning curve scanning in a tilted visual surround. | Test that tilt tuning depends of gravity and not visual cues. |
| Experiment 1-L1 |  | Free motion in arena. | Measure azimuth tuning using traditional method.<br>Control for tuning stability (by comparing with Experiment 1-L0,2). |
| Experiment 1-D |  | Free motion in arena, in darkness. | Test that azimuth tuning is maintained in darkness. |
| Experiment 1-L2 |  | Free motion in arena. | Measure azimuth tuning using traditional method.<br>Control for tuning stability (by comparing with Experiment 1-L0,1). |

**Supplemental Movie 1: Volumetric representation of 3D head orientation**. Head tilt relative to vertical is represented on a plane (labelled “2D Tilt”; see **Suppl. Fig. 5**) and head azimuth is represented on a third (unfolded) axis to form a 3D volume. The animation represents a mouse head placed at 5 different tilt orientations (upright, 60° LED, 90° ND and NU, 150° LED) and moved though all possible azimuths. The center of the head is placed at the corresponding 3D position in the volumetric plot. The movements that change azimuth without changing head tilt, correspond to rotations around an earth-vertical axis, as represented by red arrows.

**Supplemental Movie 2: 3D tuning curve of the example neuron in Fig. 1g**. The tuning curve of Fig. 1g is shown as animation sweeping through tilt angles from 0 to 60°. The lower right panel shows the average azimuth tuning curve at each tilt angle, computed by averaging across all tilt orientations. The cell is tuned to tilt, with a preferred orientation at 48° ND, but not to azimuth.

**Supplemental Movie 3: Three-dimensional passive re-orientation protocol in rotator (Fig. 1j; see also Suppl. Fig. 6) and response of a tilt-tuned RSC cell**. Upper left panel: an animated model of the 3D rotator (the rotator is ~2.4 m height; the size of the mouse head is exaggerated) displays its position recorded in real-time using potentiometers during Experiment 3-L. Lower panel: 3D head orientation is shown in volumetric space. Instantaneous head orientation is represented by a black circle; the travelled trajectory by a grey line, and recorded spikes by red dots. The trajectory is also projected on a tilt plane (right panel). A clock shows the time elapsed since the beginning of the recording. The first 10s of recording are shown in real time and then the movie accelerates to span the entire experiment. Spikes clearly cluster in a space that corresponds to ND tilt between 120 and 150°, independently of azimuth. The end of the movie (starting at t=50s) shows the complete tuning curve as a color map. The animation sweeps through all tilt angles from 0 to 180° to allow visualizing the entire 3D volume. The cell is tuned to tilt, with a preferred orientation at 135° ND, but not to azimuth.

**Supplemental Movie 4: 3D tuning curve of the example neuron in Fig. 1k (conjunctive cell**). The movie pauses at a tilt angle of 10° to show azimuth tuning, which is strongest in the vicinity of upright. The cell is also tuned to tilt, with a peak response in NU orientation. Same format as in **Suppl. Movie 2**.

**Supplemental Movie 5: 3D tuning curve of the example neuron in Fig. 1I (tilt-only cell**). Same format as in **Suppl. Movie 2**.

**Supplemental Movie 6: Azimuth tuning of an example neuron in YO, EH and TA frames**. Upper left: animated model of the rotator during Experiment 3-L. Right panels: head tilt (α) versus azimuth computed in a yaw-only frame (upper panel), earth-horizontal frame (middle) or tilted azimuth frame (bottom). The tuning curves of an example conjunctive ADN cell, obtained by averaging the 3D tuning curve across all tilt orientations (γ), are shown as a color map. As the animation runs, the position of the head and the spikes emitted by the cells are indicated in each panel. The position is identical in the 3 frames during the first few seconds of the experiment, because the head is about upright where all frames are identical. As soon as the head tilts beyond 90°, the 3 frames diverge. When the head comes back to upright orientation, the position is again identical in the EH and TA frame because these frames maintain allocentric invariance. In contrast, azimuth in the yaw-only frame can’t track allocentric head position (as shown in **Fig. 5b**). As shown by the spiking activity and the tuning curves, the cell is tuned to azimuth in the EH and TA frame. Beyond 90° tilt, where the EH and TA frame diverge most, azimuth tuning is lost in the EH frame but maintained, although weakened, in the TA frame. The final sequence of the movie shows the cell’s full 3D tuning curve. Same format as in **Suppl. Movie 2**.

**Supplemental Movie 7: 3D tuning curve of the example azimuth-only neuron in Fig. 5e**. The movie pauses at tilt angles of 30° and 110°. As shown by the azimuth tuning curve (lower right panel), azimuth tuning fades away when tilt angle increases. Same format as in **Suppl. Movie 2**.

**Supplemental Movie 8: 3D tuning curve of the example conjunctive neuron in Fig. 6b**. The neuron is tuned to tilt with a preferred orientation at 42° ND, and to azimuth with a PD of −27°. Consequently, the cell fires maximally at 3D orientations centered on these coordinates. Same format as in **Suppl. Movie 2**.

**Supplemental Movie 9: 3D tuning curve of the example conjunctive neuron in Fig. 6c; Suppl. Fig. 13**. The neuron is tuned to tilt with a preferred orientation at 105° ND, and to azimuth with a PD of 5°. Although azimuth tuning is clear in the lowest portion of the tuning curve, it decreases with tilt angle and is therefore minimal at 105°, where the cell appears to be only tuned to tilt. As a consequence, the cell seems to alternate from azimuth tuning when the head is close to upright to tilt tuning at large tilt angles. Same format as in **Suppl. Movie 2**.

**Supplemental Figure 1:**
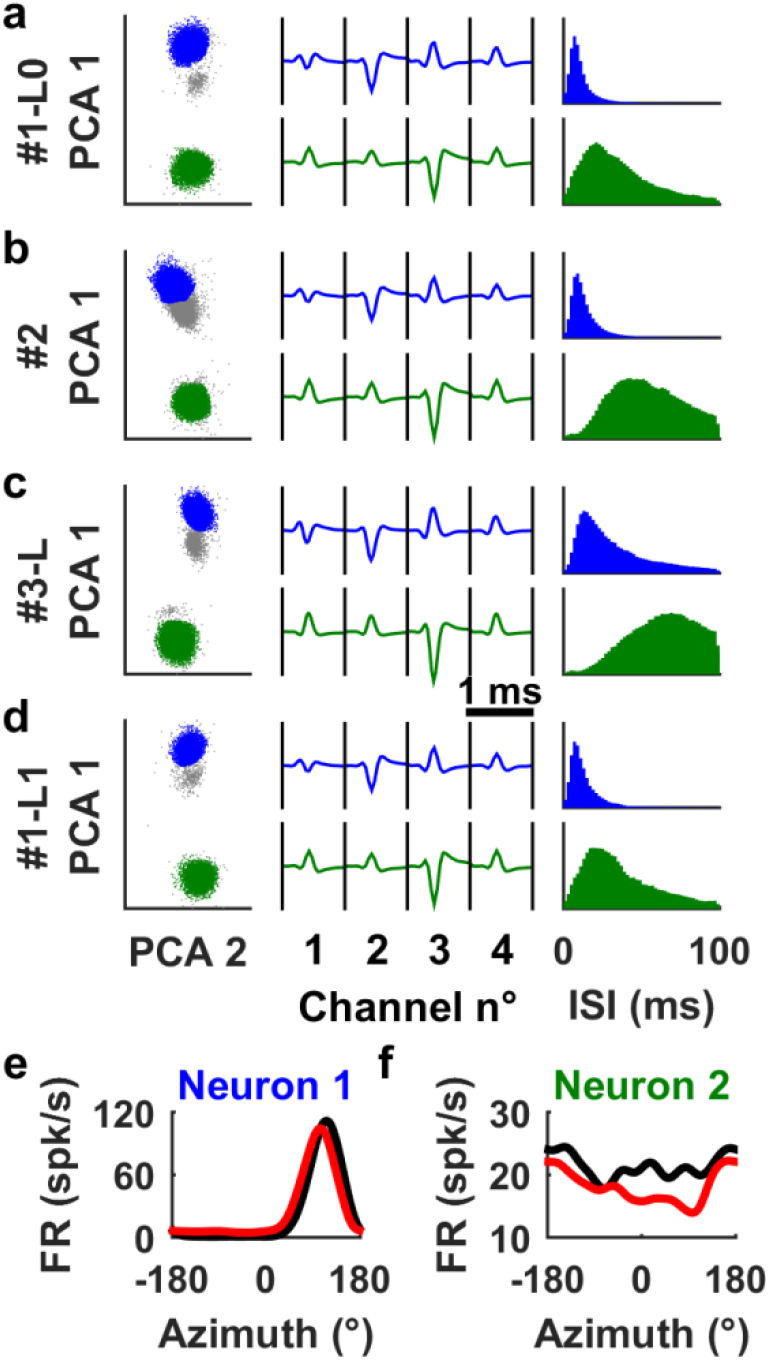
Recording stability across sessions. **(a-d)** Two neurons (color-coded in blue and green; animal H68M, CIN) were recorded simultaneously during Experiment 1-L0 (a), Experiment 2 (b), Experiment 3-L (c) and Experiment 1-L1 (d). Left panels: spikes were extracted by manual clustering based on the maximum (peak) and minimum (valley) voltage of the spikes, combined with factor analysis (the first two components, PCA 1 and PCA 2, are shown here). When represented on a 2D plot, the spikes form two clearly distinct clusters (blue and green). Grey dots represent unsorted events that result from noise and background activity. Middle panels: average spike waveforms across all channels. Right panels: the Inter-spike interval (ISI) histograms are conserved across Experiment 1-L0 and 1-L1 (a, d) but shift rightward during Experiment 2 (for neuron 2) and Experiment 3-L for both neurons. This shift reflects the reduction of average firing rate during Experiments 2 and 3 due to the general attenuation of neuronal firing in the rotator (see **Suppl. Fig. 8a,b**). **(e-f)** Azimuth tuning curves of the neurons, recorded in the initial (black) and second (red) freely moving session, i.e. before and after Experiments 2 and 3-L, are similar. This comparison serves as confirmation of recoding stability across sessions (see also **Suppl. Fig. 3**).

**Supplemental Figure 2:**
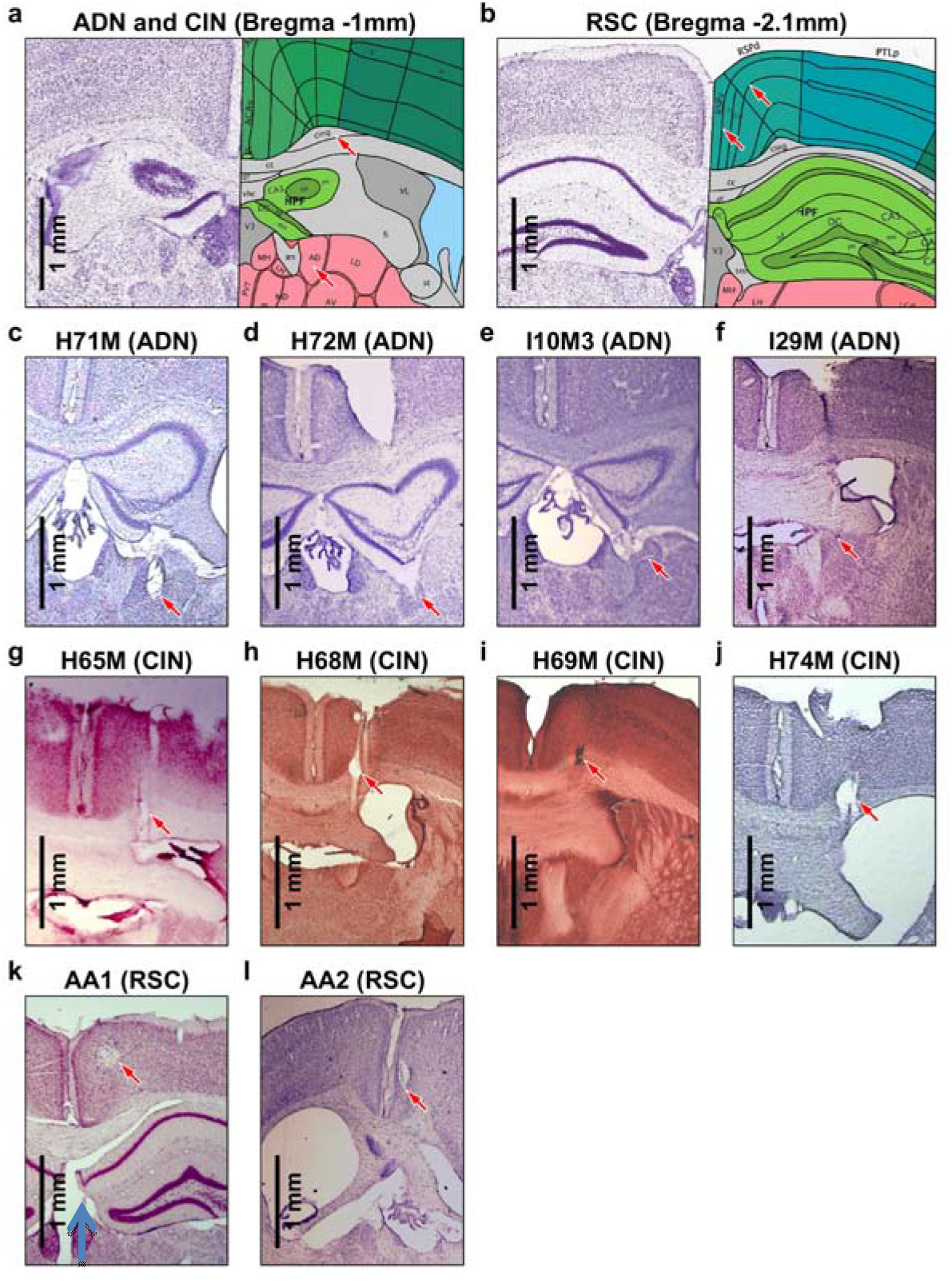
Histological localization of neuronal recordings. **(a,b)** Annotated histology slides from the Allen Mouse Brain Atlas (Lein et al., 2007; http://mouse.brain-map.org/static/atlas). The location of the sections relative to Bregma are indicated in the title. Red arrows indicate the ADN (labelled ΆD’), CIN (labelled ‘cing’) in (a), the I-IIth layer of the granular RSC (labelled RSCv; left arrow) and the V^th^ layer of the dysgranular RSC (RSCd; right arrow) in (b). Note that the ADN extends from Bregma −0.4mm to Bregma −1.1mm. Nissl-stained sections of all animals included in this study are shown; tetrode tracks are indicated by arrows. **(c-f)** Recordings in the ADN. In all animals, the ADN appears as a characteristic triangular-shaped and densely stained nucleus. In H71M, H72M and I10M3, the ADN appears below the hippocampus, i.e. more caudal than usually indicated in brain atlases (Praxinos and Franklin 2004, Lein et al. 2007). However, we confirmed that the nucleus marked by an arrow is indeed the ADN in each mouse by examining all microscopic sections and locating the anterior extremity of the thalamus as well as the anterior and posterior extent of the ADN. (**g-j**) Recordings in the cingulum fiber bundle. **(k,l)** Recordings in the RSC (AA1: dysgranular RSC, likely layer V; AA2: granular RSC; layer indeterminate).

**Supplemental Figure 3:**
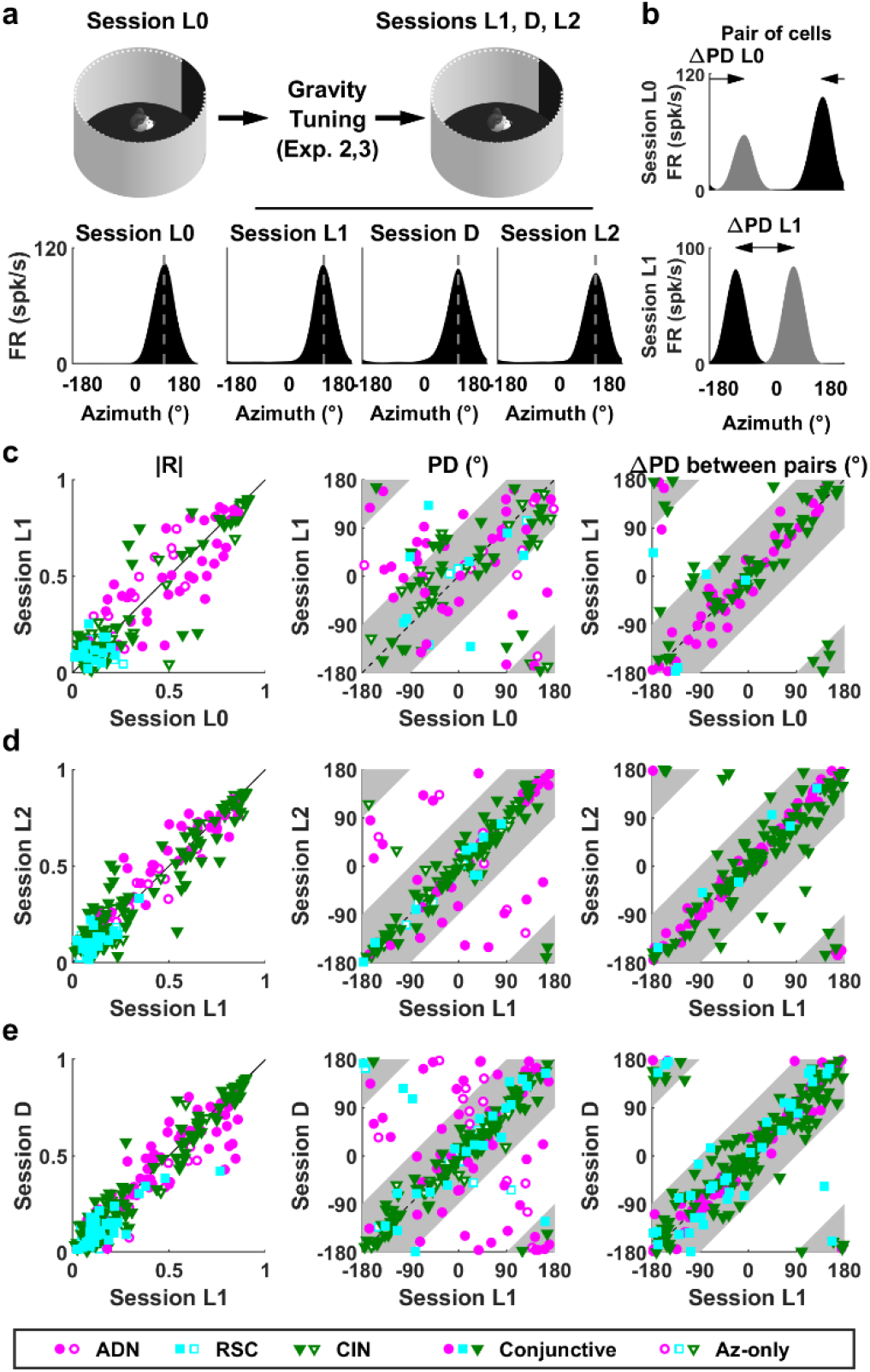
Response properties of Azimuth-tuned cells during unrestrained motion in a traditional arena (Experiment 1). As summarized in this analysis, azimuth-tuned cells conform to well-established properties (see e.g. Taube 2007, Peyrache et al. 2015). **(a)** Illustration of the sequence of recordings and example cell. At the beginning of an experimental day, azimuth tuning is recorded in light as the mouse forages freely in a circular arena (session L0). The mouse is then transferred to the platform and rotator to characterize its 3D tuning (Exp. 2-3; see **Suppl. Table 2** and **Methods**), upon completion it is returned to the freely moving arena and azimuth tuning is measured in light again (session L1), then in darkness (D), then again in light (L2). An example azimuth-tuned cell with stable preferred direction (PD) in all sessions in the arena (L0, L1, D, L2). **(b)** Two simultaneously recoded cells (grey and black tuning curves) that changed PDs between sessions L0 and L1 (see also other mouse studies: e.g., Yoder et al. 2009). Importantly, both cells shift together, such that the difference angle between their PD (ΔPD) remains constant. Thus, comparing ΔPD across cell pairs allows testing whether azimuth-tuned cells form a coherent neuronal compass even when this compass drifts from one session to another. **(c)** Azimuth response stability between sessions L0 and L1. Left: There is no significant difference in tuning strength (Mean vector length |R|: signed rank tests, p>0.5 for all groups; Bonferroni correction applied; data from all cells significantly tuned to azimuth in at least one session; n = 54 ADN; 33 RSC; 83 CIN). Middle: Comparison between the PD of individual cells. Only HD cells significantly tuned (p<0.01) in both sessions are included. PDs of a small subpopulation may drift between session L0 and L1 (PD shift > 90° in 14/46 ADN; 3/12 RSC; 7/56 CIN cells, i.e. 21% cells total). Grey bands represent sectors where the PDs shift by less than 90°. Right: ΔPD between pairs of simultaneously recorded cells. Only cells significantly tuned (p<0.01) in both sessions are included. PD differences are stable (<90° shift) in 93% (45/46 ADN; 4/5 RSC; 50/55 CIN) of cells pairs. Thus, although PD may shift between L0 and L1, the PD of all cells tend to shift together, in line with predictions of an attractor network (Peyrache et al. 2015). **(d)** Azimuth response stability between sessions L1 and L2 (same legend as in c). There is no significant difference in tuning strength (p=0.03 for ADN, p>0.5 for other groups; n=55 ADN; 33 RSC; 135 CIN). Only 10% of cells (15/48 ADN; 0/15 RSC; 1/92 CIN) drift more than 90°. PD differences are stable (<90° shift) in 97% (61/61 ADN; 6/6 RSC; 102/108 CIN) of cell pairs. Thus, PD are more stable between L1 and L2 compared to L0 and L1, likely because of the shorter time interval between L1 and L2 and/or the use of 3D stimuli in-between sessions L0 and L1. **(e)** Azimuth response stability between sessions L1 and D (same legend as in c,d). Tuning strength is slightly attenuated in darkness in RSC (linear regression slope=0.71, p < 10^−5^; n=70) but not in other areas (ADN: n=70, p = 0.14; CIN: n=143; p = 0.3). Only 14% (25/63 ADN; 4/37 RSC; 0/113 CIN) of PDs drift more than 90°. PD differences are stable (<90° shift) in 99% (84/84 ADN; 38/39 RSC;146/148 CIN) of cell pairs.

**Supplemental Figure 4:**
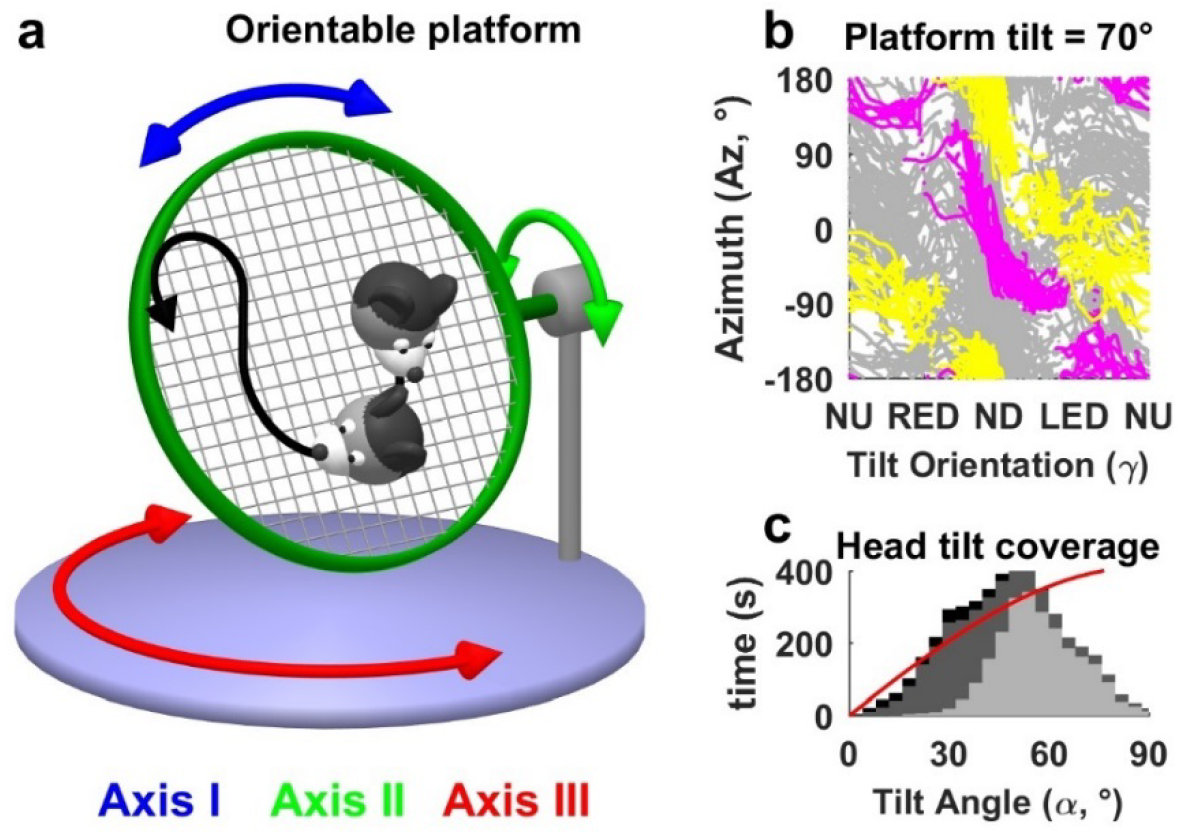
Freely moving setup and protocol for measuring 3D tuning in unrestrained, exploring mice. **(a)** Illustration of the 3D orientable platform setup: mice walk freely (black arrow) on a meshed platform that can rotate around 3 axes (blue, green, red). Walking on the platform changes tilt orientation (γ; see **Suppl. Fig. 5**) and azimuth simultaneously. Rotating the base (Axis III) changes azimuth but not tilt orientation. Axis II is used to change tilt angle (α; see **Suppl. Fig. 5**). Recordings are performed in 5-8-minute blocks where mice walk freely while axis II-III are set to a static position. Axis I is rotated randomly in the middle of each block to prevent tilt orientation relative to gravity from being coupled to head orientation relative to local cues on the platform itself. Because of this added complexity, earth-horizontal azimuth tuning in Fig. 2a was evaluated based on data recorded in the arena. **(b)** Distribution of azimuth (Az) and tilt orientation (angle γ) in eight 5-min blocks where the platform was tilted 70° (average head tilt = 60±15°, as mice tend to partially compensate with their head). Yellow, magenta: data recorded during two 5-min blocks corresponding to different configurations (same tilt angle, but different positions of Axis III). Azimuth and tilt orientation vary together as animals walk within one block, but azimuth is offset when Axis III is rotated between blocks, thus allowing to scan the entire tilt orientation/azimuth plane. **(c)** Distribution of head tilt angle (α) in the same recording session (black/grey/light grey: data collected with the mesh tilted 0°, 45°, 70°, respectively). Red curve: distribution required for uniform sampling of tilt orientation, illustrating relatively uniform sampling up to ~60°).

**Supplemental Figure 5:**
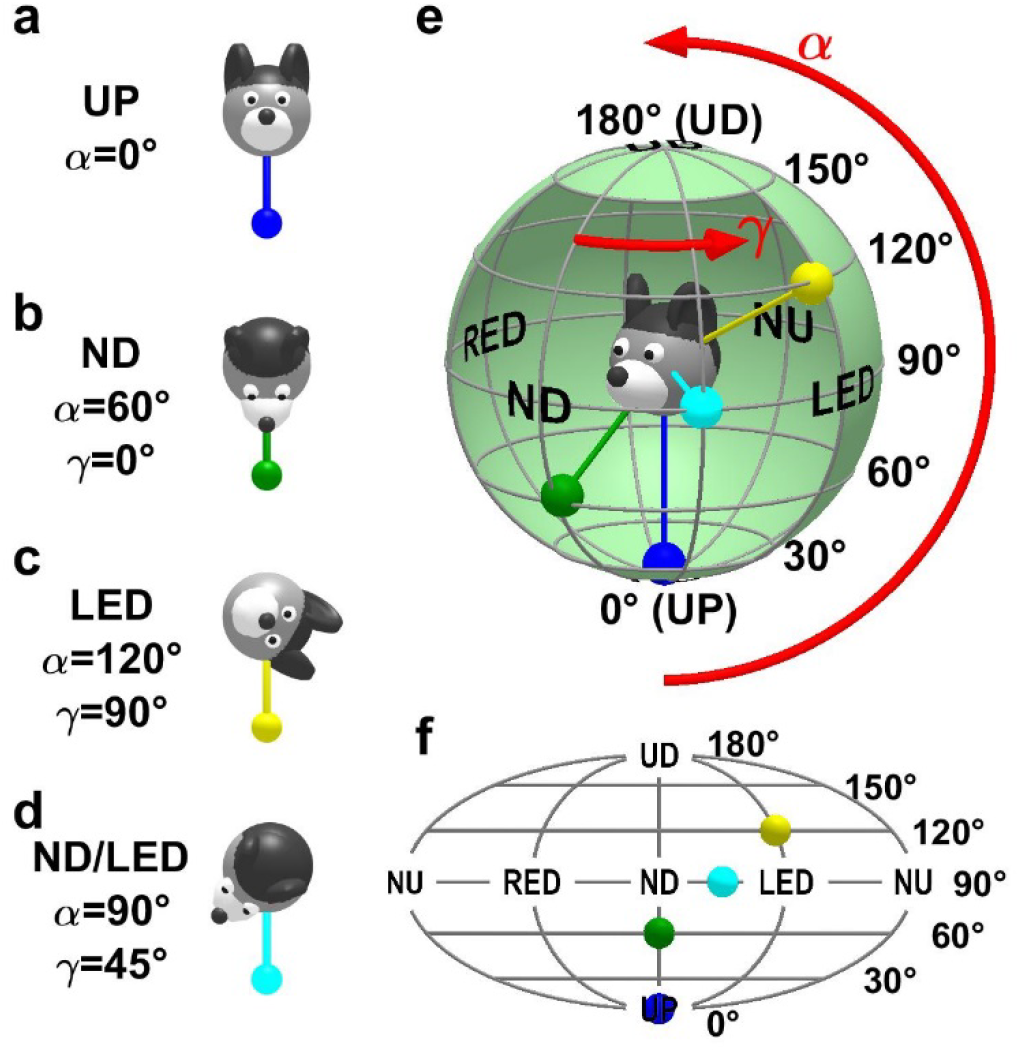
Spherical coordinate system for tilt orientation and Mollweide projection. The rationale for the coordinate system is the following: each tilt orientation relative to gravity is equivalent to, and defined as, the corresponding orientation of the gravity vector relative to the head. **(a-d)** Four example tilt orientations, expressed in a spherical coordinate system (α, γ) where α is the tilt angle and γ is tilt orientation: γ = 0° and γ = 180° correspond to nose-down (ND) and nose-up (NU) tilt; γ = 90° and γ = −90° correspond to left-ear-down (LED) and right-ear-down (RED) tilt. The colored pendulum/ball represents the gravity vector. **(e)** Spherical topology of tilt orientation. When head tilt spans all possible orientations, the tip of the gravity vector spans a sphere surrounding the head. The tilt variable α corresponds to the latitude on the sphere. Upright (UP, α=0°) and upside-down (UD, α=180°) orientations correspond to the lower and upper pole respectively. The orientation variable γ corresponds to the longitude. 90° tilt in ND, LED, NU and RED orientations are marked. **(f)** Planar representation of the sphere using an equal-area Mollweide projection. The 4 tilt orientations in (a-d) are marked with color balls.

**Supplemental Figure 6:**
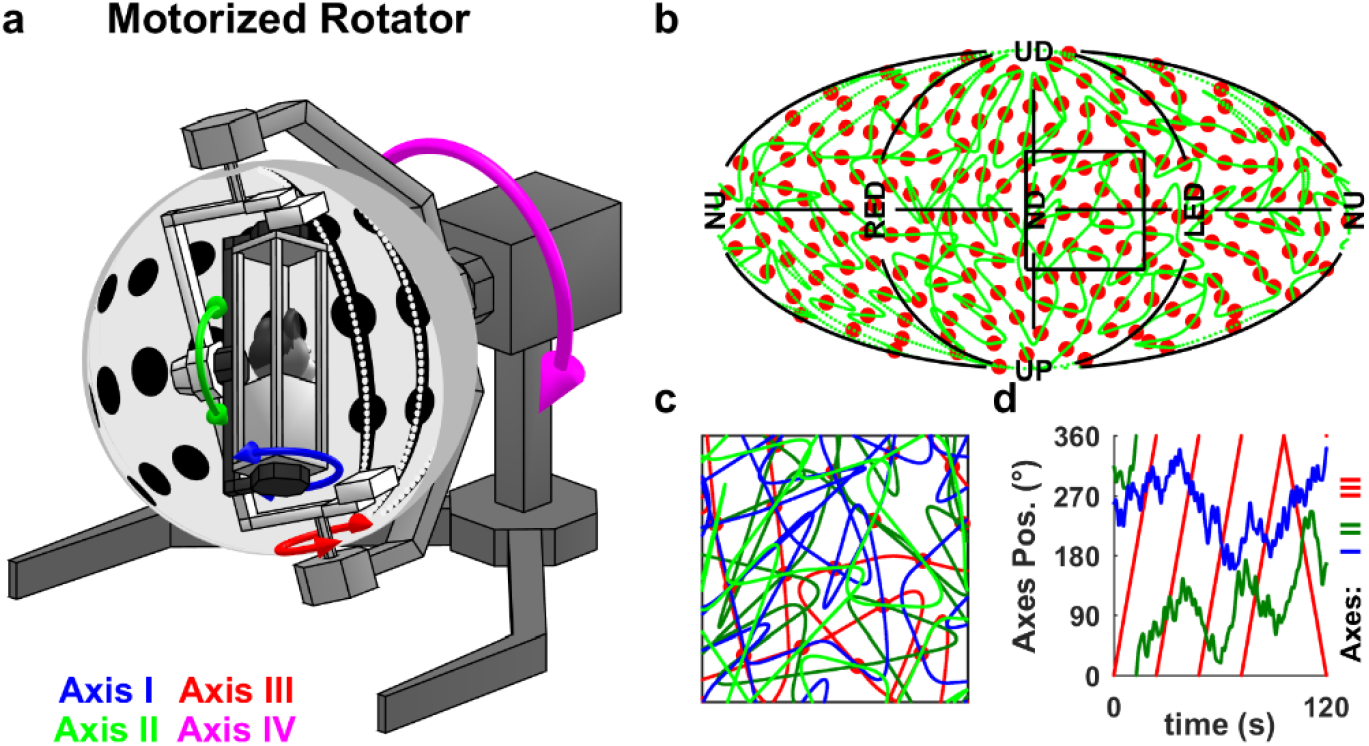
Motorized rotator setup and protocol for measuring 3D tuning in restrained mice. **(a)** Illustration of the motorized rotator; the 4-rotation axes are indicated by colored arrows. **(b)** Pseudo-random trajectory (green curve) used to measure tilt tuning. The trajectory visits 200 uniformly distributed tilt positions (red dots). The full protocol scans the entire tilt space 8 times by running through 4 distinct trajectories, each of which is ran twice in opposite directions. **(c)** Detail of the highlighted square in (b), with 4 distinct trajectories. **(d)** Position of the rotator’s 3 inner axes during a 2 min segment of the motion. Axes I (inner yaw, blue) and II (middle, pitch/roll, green) are used to manipulate 2D head tilt relative to vertical, while axis III (outer yaw, red) is used to continuously vary azimuth. Axis IV is used to tilt the setup in Experiment 3-T (**Suppl. Table 1**).

**Supplemental Figure 7:**
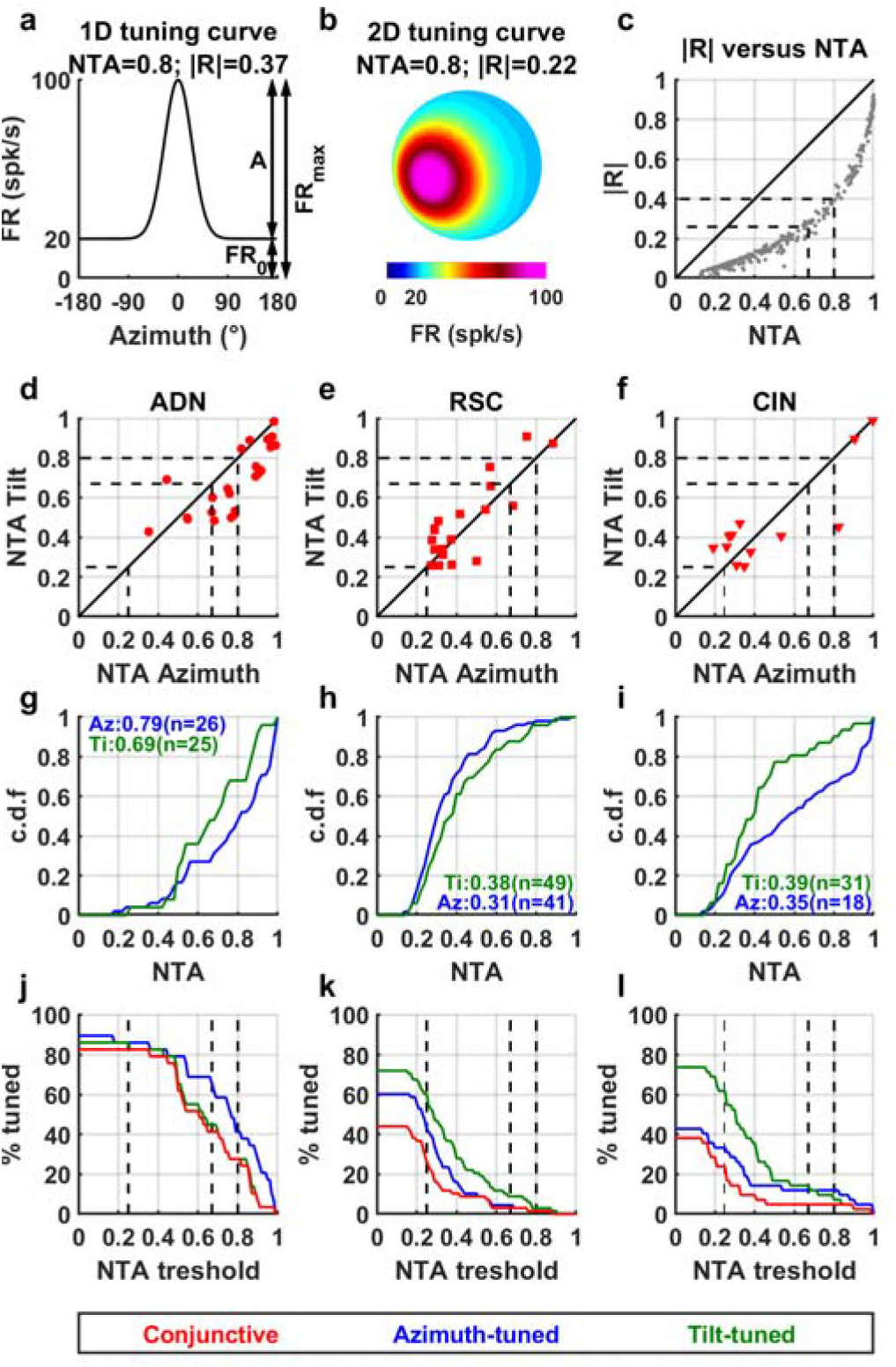
Normalized tuning amplitude vs. mean vector length for quantifying tuning strength for fair comparison between tilt- and azimuth-tuned cells. **(a)** We define a measure of tuning strength called *normalized tuning amplitude (NTA)*, illustrated here for a 1D tuning curve. We decompose firing rate into a baseline firing FR_0_ and a modulation amplitude A. The peak firing rate is FR_max_=FR_0_+A. Normalized tuning amplitude is defined as NTA=A/FR_max_. The normalized tuning amplitude of this example is NTA = 0.8; and the mean vector length is |R| =0.38. **(b)** Illustration of a 2D tuning curve on a sphere, used to model tilt tuning. This distribution has identical baseline, amplitude and standard deviation as in (a) and therefore, the normalized tuning amplitude is also NTA = 0.8. However, the mean vector length is |R| =0.22; i.e. lower than in (a). This is because the area of baseline firing on a sphere (cyan in b) amounts to a larger portion of the tuning curve than in a 1D tuning curve, resulting in a lower |R| value. Therefore, the mean vector length is inappropriate for comparing azimuth (1D) and tilt (2D) tuning. **(c)** Comparison of the mean vector length (|R|) and normalized tuning amplitude for azimuth tuning in Experiment 1-L (**Suppl. Table 2;** data from all significantly-tuned cells). Although the scaling between the two measures is non-linear, there is a clear relationship between them, indicating that they provide the same information. *Therefore, the normalized tuning amplitude is as suitable as the mean vector for quantifying tuning strength, with the advantage that it allows for a fair comparison between 1D and 2D tuning curves*. **(d-f)** Comparison of the normalized tuning amplitude of azimuth and tilt tuning for all conjunctive cells (paired data from Experiment 1-L and Experiment 2; **Suppl. Table 2**; only cells tested in both experimental protocols and significantly tuned to both Az and tilt have been included; ADN: n=24; RSC: n=19; CIN: n=12). Azimuth tuning is slightly higher than tilt tuning in ADN (median A_N_ = 0.79 versus 0.7, p = 0.002, signed rank test) but not in RSC (0.37 versus 0.44, p = 0.2) nor in CIN (0.34 versus 0.41, p = 0.8). Previous studies have classified neurons as HD cells when |R|>0.26 (Jacob et al. 2017) or |R|>0.4 (Yoder et al. 2009). We infer from the population response in (c) that these values correspond to NTA≈0.67 and NTA≈0.8, respectively. In the present study, we classified cells as azimuth or tilt-tuned using statistical criteria: if tuning passed a shuffling test (at p<0.01), as long as NTA>0.25 (to exclude low-modulating cells). The three thresholds (NTA=0.25, 0.67, 0.8) are indicated by vertical dashed lines. Although the statistical criterion we used was more inclusive compared to the fixed threshold of previous studies, the 3D tuning properties described here are found in those cells that pass the more restrictive criteria of previous studies, as detailed in subsequent panels. **(g-i)** Cumulative distribution of NTA for all azimuth-tuned (blue) and tilt-tuned (green) cells that passed the shuffling test (not only conjunctive cells as in panels d-f). Median values of NTA and number of cells are indicated in the panels. The median values of tilt and azimuth tuning are comparable in all regions. **(j-l)** Percentage of cells that would be classified as azimuth-tuned (blue), tilt-tuned (green) and conjunctive (red) by passing the shuffling test (p<0.01) and exceeding a variable NTA threshold, expressed as a function of that threshold, but including all recorded cells in both the arena and platform setups. Note that the threshold value of NTA =0.25 used in the present study allows for a large fraction of cells to be classified as azimuth-tuned, tilt-tuned or conjunctive (e.g. 86% azimuth-tuned and 86% tilt-tuned in ADN). Using a more stringent threshold (e.g. NTA=0.8) to select cells with very vigorous responses, we find that 41% ADN cells are classified as azimuth-tuned, 28% as tilt-tuned, and 28% as conjunctive. Thus, even when a stringent criterion is used, over a fourth of ADN cells exhibit large 3D responses. A sizeable, but lower, fraction of CIN cells (12% azimuth-tuned, 7% tilt-tuned, 5% conjunctive) exhibit similar strong responses. In contrast, only a minority (<3%) of RSC cells pass this threshold, indicating that significant HD responses exist in RSC but are generally weaker than in ADN, which is a known property already (Chen et al. 1994a). Importantly, the proportion of tilt-tuned cells is typically higher that Az-tuned cells in all areas, regardless of NTA threshold (green vs. green curves).

**Supplemental Figure 8:**
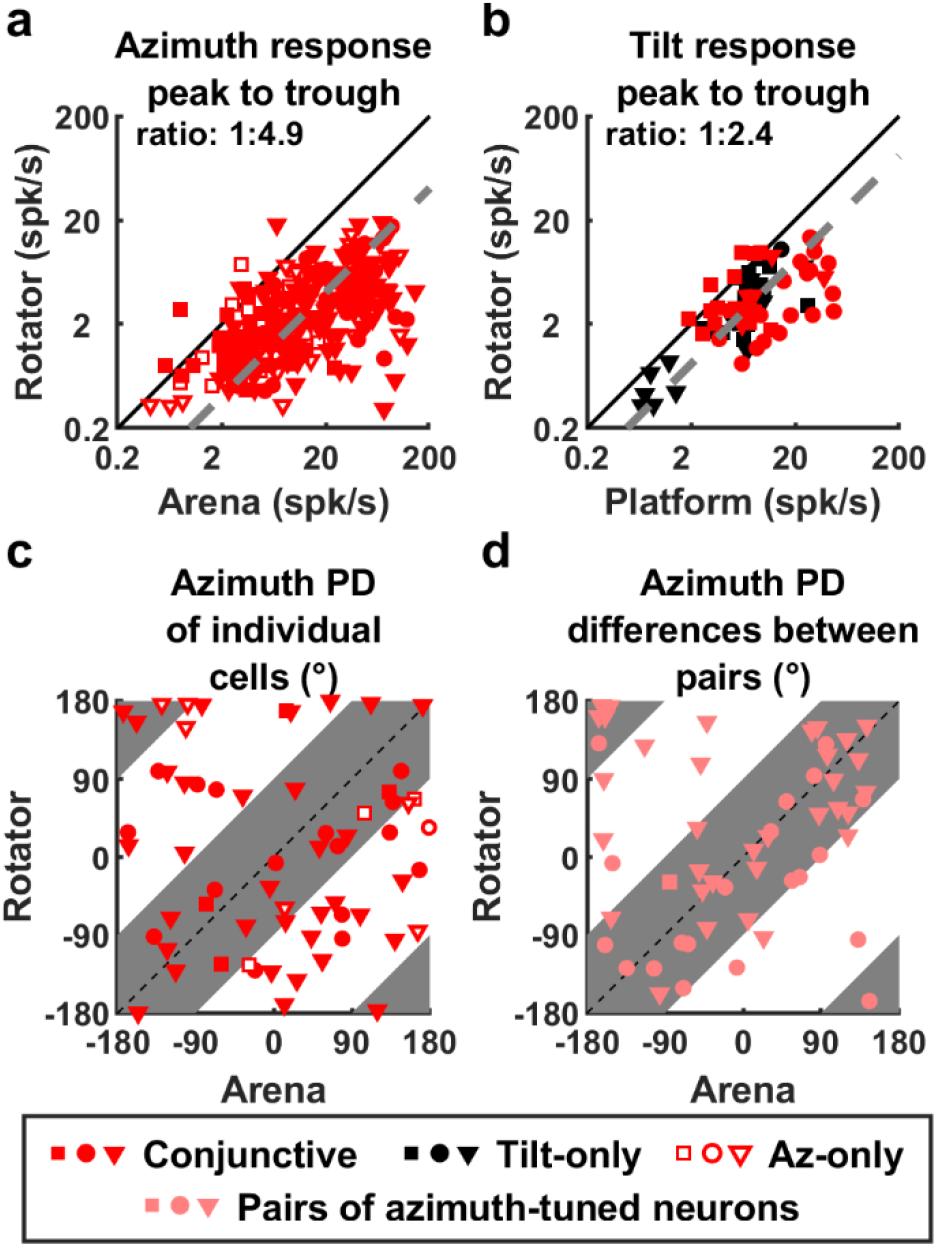
Comparison of azimuth and tilt responses when moving freely and restrained. **(a)** Comparison of azimuth response amplitude when moving freely in the arena (Experiment 1-L0,l,2; **Suppl. Table 2**) and when restrained in the rotator (Experiment 3-L, considering data for tilt < 45°). Data from cells significantly modulated to azimuth in Experiment 1 and recorded during Experiment 3-L (n=267). Responses were largely attenuated when mice were restrained (median amplitude ratio= 1:4.9 [1:4.4 - 1:6.3] Cl; grey). **(b)** Comparison of tilt response amplitude when moving freely on the platform (Experiment 2; **Suppl. Table 2**) and when restrained in the rotator (Experiment 3-L; re-analyzed based on data when head tilt < 60° to match the range of tilt sampled in Experiment 2). Data from cells significantly modulated to tilt when moving freely and recorded during Experiment 3-L (n = 70). Responses were attenuated when mice were restrained (median amplitude ratio= 1:2.4 [1:1.9 1:2.8] Cl; grey). Together, panels (a, b) indicate that restraining mice leads to an attenuation of both azimuth and tilt response amplitude. Azimuth tuning is attenuated to a larger extent than tilt tuning, which is the reason tilt tuning curves can be reliably measured in the rotator in most neurons, but azimuth tuning curves only in a minority of neurons (**Fig. 2c**). M) Inconsistency in azimuth PDs, but consistency of difference in azimuth PD between pairs of cells across setups. The arena and the rotator are different setups located in separate rooms and don’t share a common azimuth reference. Therefore, the azimuth compass may anchor to a priori random orientations in each setup, which would cause the PD of individual cells to vary randomly. Accordingly, PDs shift by more than 90° in 29/63 cells (panel c, p=0.6, Binomial test). However, if azimuth-tuned cells are part of a neural compass, then the PD of simultaneously recorded cells should remain anchored one relative to the other. Therefore, the difference in PD between pair of cells should be identical in the arena and in the rotator. Indeed, PD differences in the arena vs. rotator were significantly conserved in panel d (within 90° in 44/54 pairs of cells, p<10^−5^, Binomial test). Note, however, that unlike the azimuth compass that anchors to visual landmarks, a tilt compass anchored to gravity (**Fig. 1a**), should have identical PDs on the platform and in the rotator. This is indeed the case, as shown in the pixel-by-pixel correlation of the fitted tilt tuning curves up to 60° tilt that could be tested in both setups (**Fig. 2d**; We used the correlation analysis for tilt tuning, because the PD of most cells can’t be measured on the platform, which is restricted to 60° tilt. Also note that together, panels a-d and **Fig. 2** indicate that the spatial tuning characteristics of both azimuth and tilt tuning are conserved across free locomotion and restrained, passive motion, and that, other than the smaller response magnitude, the 3D responses measured in the rotator are representative of the neurons’ natural responses.

**Supplemental Figure 9:**
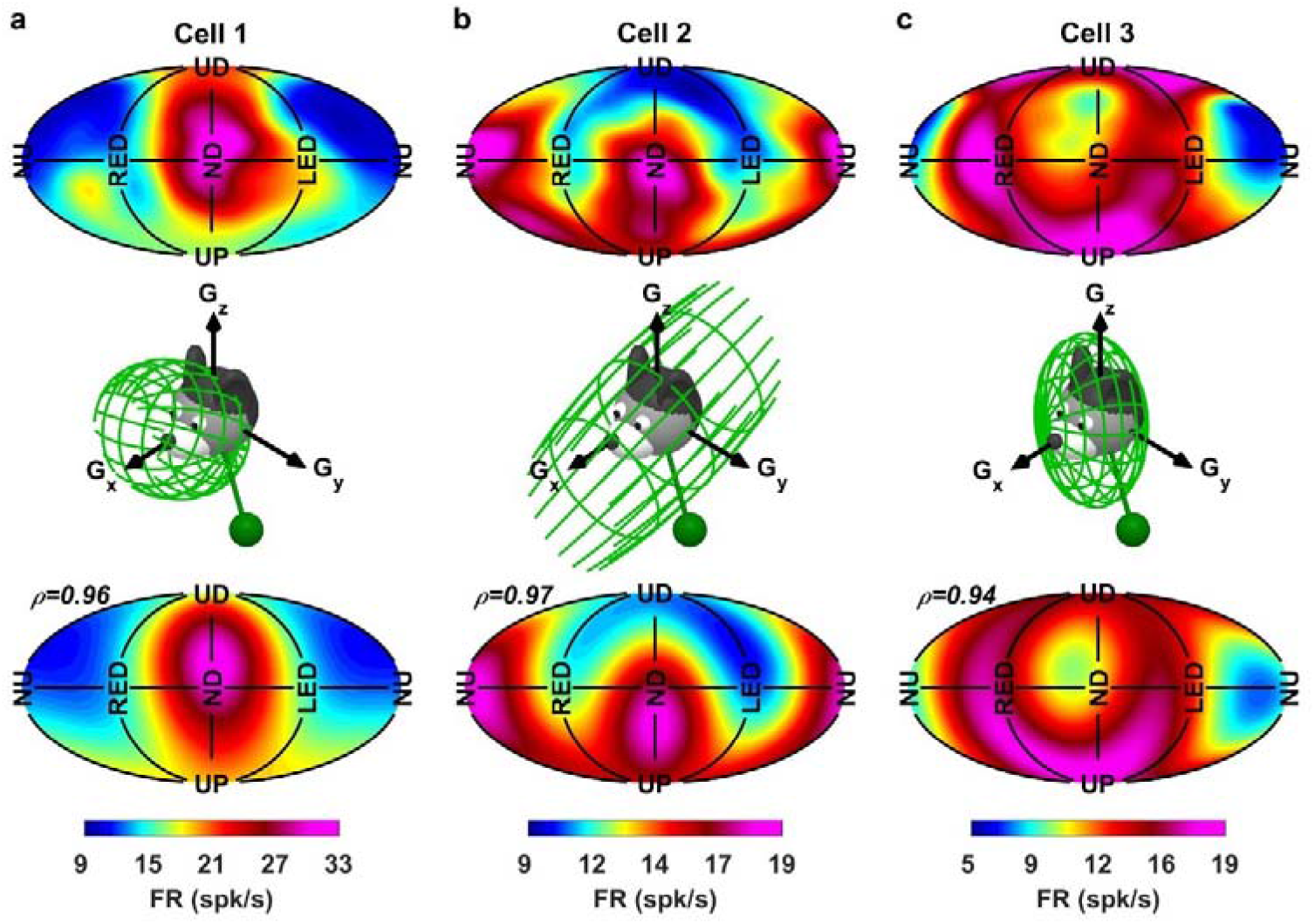
Example tilt tuning curves fitted with a Gaussian function. **(a)** Example cell with a single hill of activity (representative of the majority of cells). Top panel: planar representation of the tilt tuning curve (PD close to ND). Middle panel: illustration of the Gaussian fitting. Bottom panel: planar representation of the fitted firing rate. The correlation between the measured and fitted firing rate (top and bottom panel) was p=0.96. **(b)** Example cell with a bimodal response. The cell responds preferentially in NU and ND orientations. Middle panel: When gravity is expressed in Cartesian coordinates, NU and ND orientations correspond to the gravity vector being aligned with the naso-occipital axis (G_X_), and therefore G_Y_≈0 and G_Z_≈0. A 3D Gaussian with a large standard deviation along G_X_ but small standard deviation along G_Y_ and G_Z_ will have a large value at all positions close to the G_X_ axis and therefore resemble the cell’s tuning curve. Accordingly, the cell’s firing was fitted with an elongated 3D Gaussian whose long axis was aligned with G_X_ (shown as a green wireframe ellipse that corresponds to one std. dev.). The correlation between the measured and fitted firing rate (top and bottom panel) was ρ=0.97. **(c)** Example cell with a tuning curve exhibiting a ring-shaped hill of activity (top panel), responding preferentially in UP, UP, RED and LED but not in ND or NU. Middle panel: In Cartesian coordinates, the cell’s response corresponds to gravity being aligned with the G_Y_, G_Z_ plane. Accordingly, the cell’s response was fitted with a flat, pancake-shaped 3D Gaussian that has a large value along the G_Y_, G_Z_, but small values along the G_X_ axis (ρ=0.94).

**Supplemental Figure 10:**
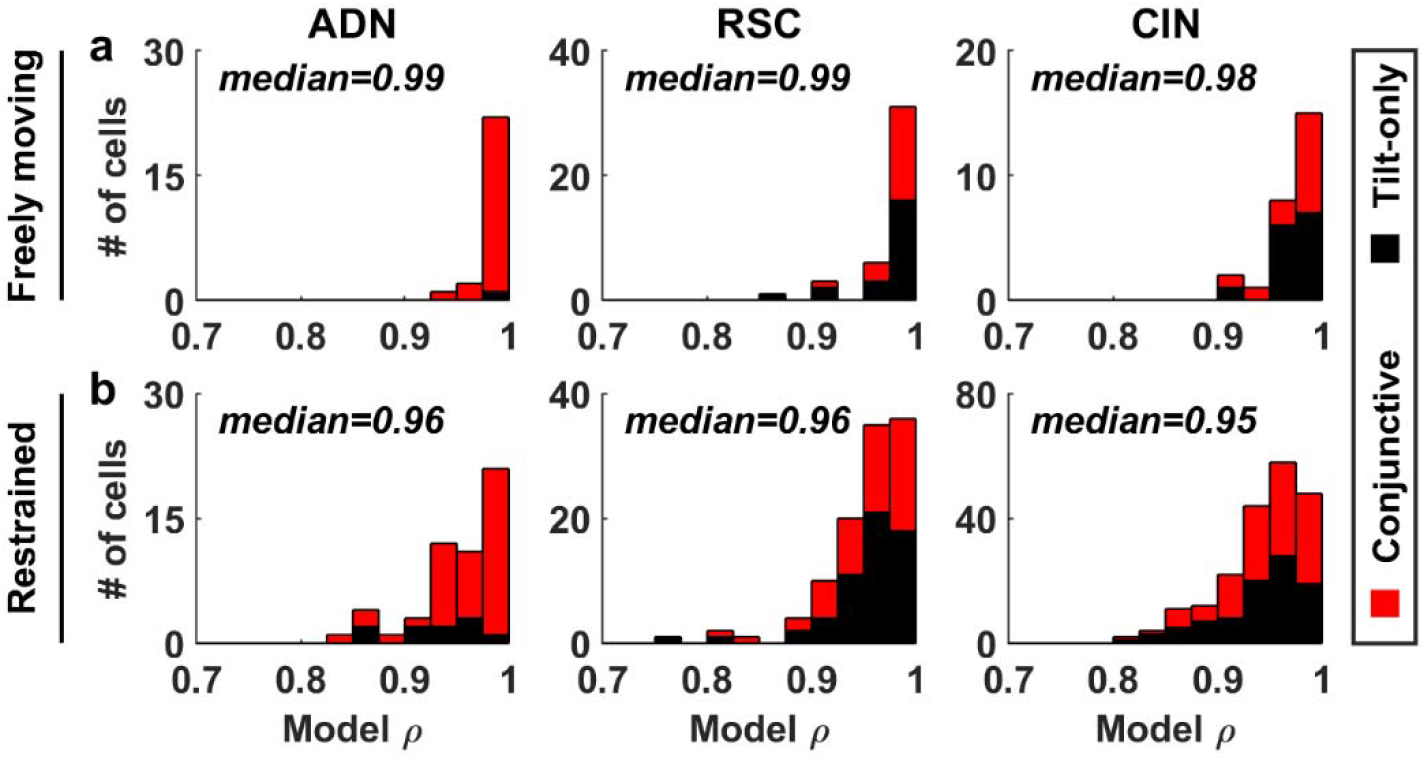
Summary of goodness of fit (coefficient of correlation, ρ) of Gaussian functions to 3D tuning curves, separated by brain area. Red/black: conjunctive and tilt-only cells. **(a)** Gaussian fit to the tilt tuning curves measured when moving freely. The correlation doesn’t vary between regions or between conjunctive and tilt-only cells (2-way ANOVA: p=0.33 and p=0.45 respectively). **(b)** Gaussian fits to the tilt tuning curves measured when mice are restrained in the rotator. The correlation doesn’t vary between regions or between conjunctive and tilt-only cells (2-way ANOVA: p=0.13 and p=0.26 respectively). The model’s correlation is higher (median: 0.99 versus 0.96, p<10^−10^, Wilcoxon rank sum test), likely because of the small response amplitude when restrained.

**Supplemental Figure 11:**
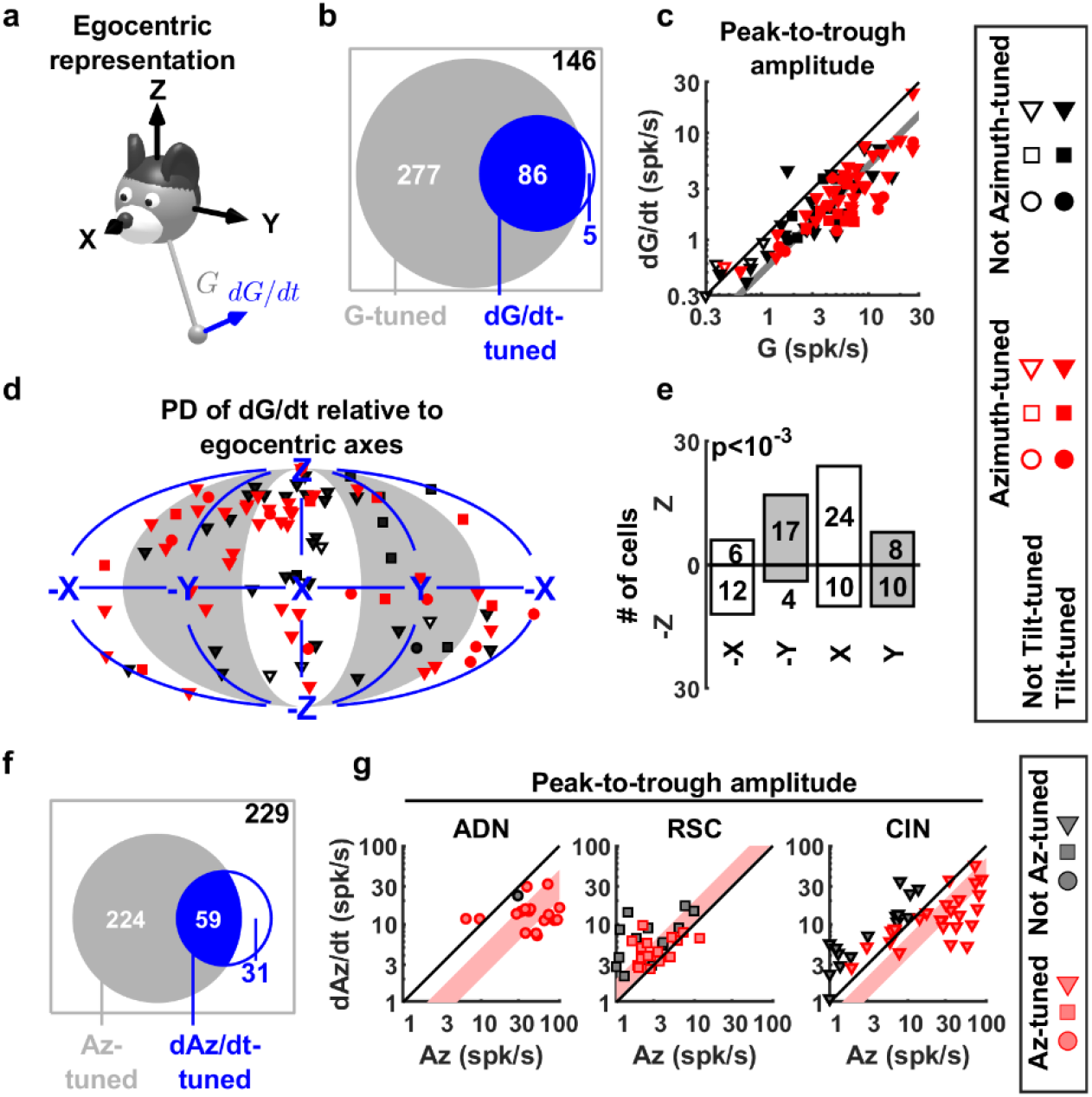
Responses to tilt and azimuth velocity. We used identical criteria to assess whether cells were significantly tuned to azimuth velocity (dAz/dt) and tilt velocity (i.e. the time derivative of gravity, dG/dt) (see Methods). **(a)** Gravity derivative was expressed in an egocentric (X,Y,Z) frame, similar to gravity, and the responses to gravity derivative was fitted with Gaussian functions (as in **Suppl. Fig. 10**). **(b)** A small percentage of cells (91/514, 18%) exhibited significant tuning to tilt velocity (data from Experiment 3-L; **Suppl. Table 2**). Furthermore, the majority (86/91; 95%) of these were also tuned to tilt. Tilt-tuned cells were more likely to be tuned to dG/dt (p<10^−7^, Chi square test). **(c)** For most cells, tilt velocity responses had a lower amplitude than tilt position responses (geometrical average ratio=0.5, [0.44 0.55] Cl; data from n=86 cells with significant tilt and tilt velocity tuning). There was no significant difference between areas (Kruskal-Wallis ANOVA, p=0.05). **(d)** Distribution of PDs for tilt velocity. ‘X’, ‘Y’, ‘Z’ indicate that cells fire preferentially when dG_X_/dt>0, dG_Y_/dt>0, and dG_Z_/dt>0, respectively. ‘-X’, ‘-Y’, ‘-Z’ indicate that cells fire preferentially when dG_X_/dt<0, dG_Y_/dt<0, and dG_Z_/dt<0, respectively. **(e)** Number of cells in all **8** quadrants of panel d. Most cells prefer dG_Z_/dt>0, and dG_X_/dt>0, i.e. when the gravity vector moves forward and upward in head coordinates, which corresponds from instance to ND pitch movements when starting from an upright condition. **(f)** A small percentage of cells (90/543,17%) was also tuned to azimuth velocity (dAz). Azimuth-tuned cells were more likely to be tuned to dAz (66% vs. 49%; p=0.005, Chi square test), (g) For cells tuned to both Az and dAz, Az velocity responses had a lower amplitude than Az position (direction) responses in ADN (median ratio: 1:3.1, [2-4.9] Cl; p < 10^−3^, signed rank test), but in contrast were slightly larger RSC (median ratio 1.5:1, [1.1-2] Cl; p < 10^−3^). The ratio in CIN was intermediate (2:1 in favor of Az responses, [1.4-2.8] Cl; p < 10^−5^). The peak-to-trough amplitude of dAz tuning curves, measured across the range of ±200°/s, had a median value of 13 spk/s ([11-15] Cl) in ADN, 5.5 spk/s; [3.7-6.6] Cl) in RSC. The distribution of responses in CIN resembled a mixture of ADN cells (with high Az and lower dAz responses) and RSC cells (with low Az and dAz responses).

**Supplemental Figure 12:**
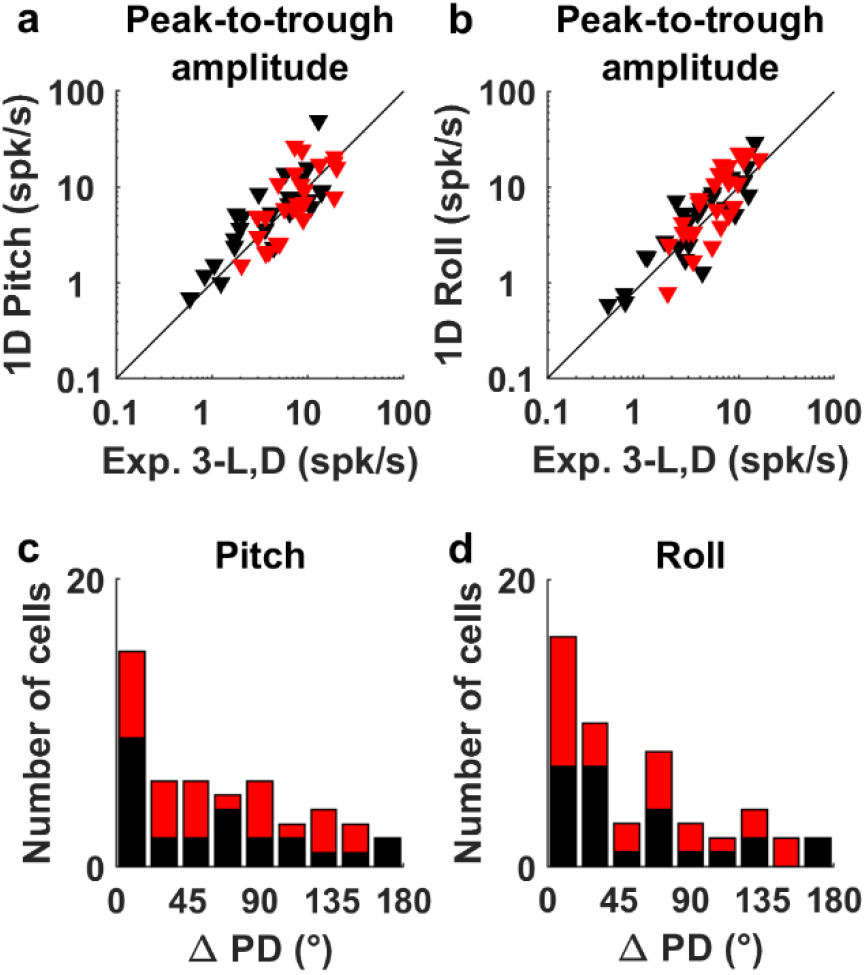
Comparison between modulation amplitude and preferred direction of tilt-tuned cells during pitch/roll rotation (Experiment 4; Suppl. Table 2) and 3D rotation (Experiment 3-L,D) (a), (b) Peak-to-trough modulation amplitude measured during pitch/roll (ordinate) vs. predicted based on tilt tuning curves measured in Experiment 3-L,D (abscissa). Amplitudes are significantly correlated (p<10^−10^ for both pitch and roll; n=50 tilt-tuned cells, data averaged across recordings in light darkness). The responses are slightly higher during single axis rotation in roll (median=5.9 vs 3.9 spk/s; p =10^−3^, signed rank test) but not pitch (median=6.4 vs 4.4 spk/s; p =0.3, signed rank test). **(c), (d)** Distributions of absolute difference in tilt preferred direction (PD) between 1D and 3D stimuli for pitch and roll planes, respectively. Both are significantly aligned with 0 (Kolmogorov-Smirnov tests to test the difference with a uniform; pitch: p < 10^−4^; roll: p = 2.10^−3^). Red symbols/bars: Azimuth-tuned cells; Black symbols/bars: Not-azimuth-tuned cells.

**Supplemental Figure 13:**
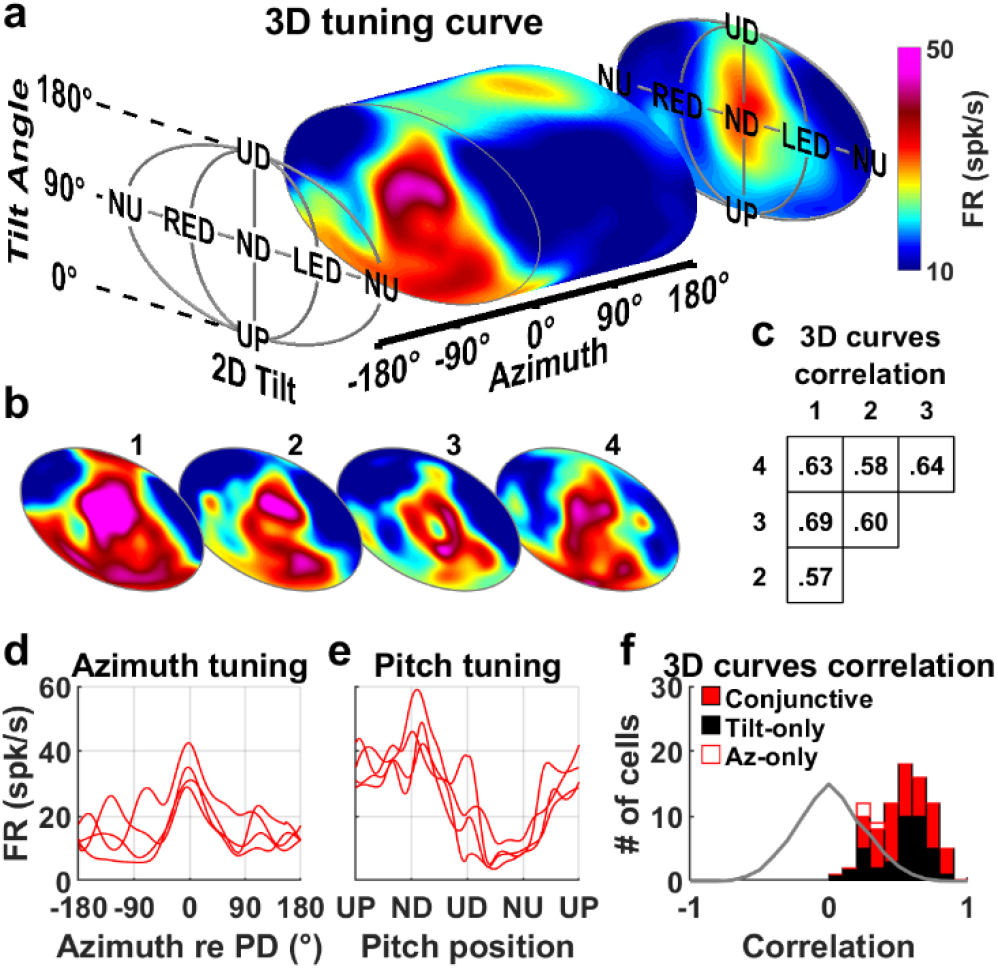
Reproducibility of 3D tuning across days. **(a)** 3D tuning curve of an example conjunctive CIN cell (same as in **Suppl. Movie 8**). A vertical section of the 3D tuning curve is shown at an azimuth of 0° that corresponds to the cell’s PD. Data averaged across all repetitions of Experiment 3-L. **(b)** Reproducibility of the tuning curve. The cell’s response was recorded 4 times during Experiment 3-L, on 4 distinct days spanning a 2-week period. 3D tuning curves were recomputed for each repetition. The same vertical section as in (a) is shown for all repetitions (labelled 1 to 4), using the same color scale. Peak firing occurs consistently in the vicinity of ND orientation. **(c)** Pixel-by-pixel correlation, across the entire 3D space, between all 4 tuning curves. Each curve is correlated with all others (p<10^−10^). The average correlation is 0.62. **(d)** Azimuth tuning curve in upright orientation extracted from the 4 3D tuning curves, and centered on each curve’s PD. **(e)** Pitch tuning curve extracted from the 3D tuning curves, at the azimuth corresponding to each curve’s PD. Analyses in panels b to e indicate that 3D responses (or responses along 1D yaw and pitch trajectories) were stable across several days in the example cell. **(f)** Distribution of the average correlation between repetitions of Experiment 3-L (as in panel c), for all tuned cells (n= 88 cells; 45 conjunctive, 39 tilt only, 3 azimuth-only). The median correlation is 0.52 ([0.49-0.58] Cl) and is similar for conjunctive and tilt-only cells (Wilcoxon rank sum test, p=0.77). Grey: expected distribution if tuning curves shift randomly between days (H0), computed by randomly shuffling the cells (Kolmogorov-Smirnov test of observed distribution versus H0: p<10^−10^). This analysis demonstrates that 3D responses are stable across days.

**Supplemental Figure 14:**
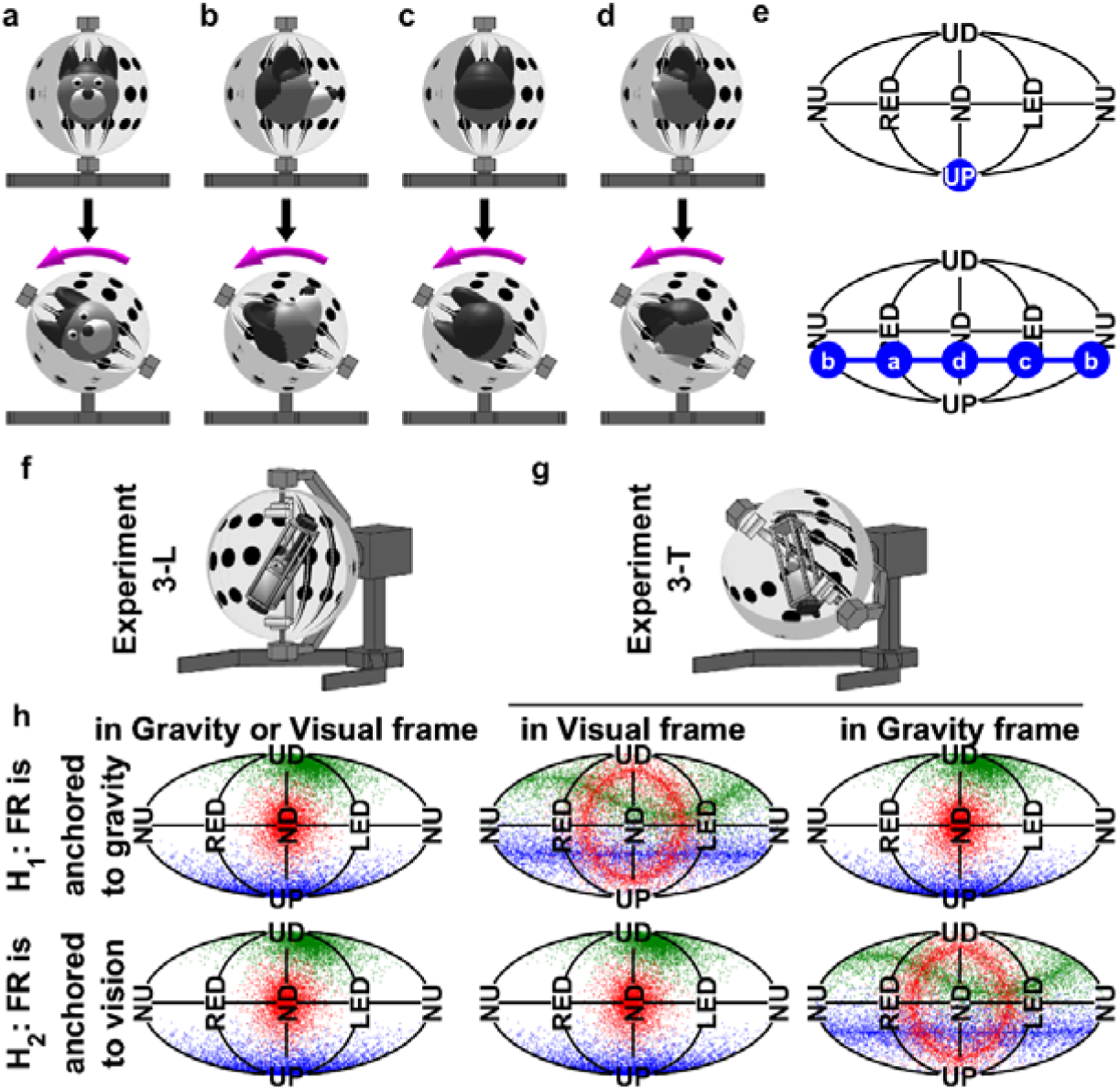
Protocol 3-T. **(a-e)** Change of head tilt resulting from rotating Axis IV. Panels a-d illustrate 4 situations where the head is initially upright and facing various direction in the horizontal plane. When Axis IV is tilted 60° (magenta arrow), the head reaches 60° in RED, NU, LED or ND, depending of its initial orientation in the horizontal plane, i.e. relative to Axis IV. **(e)** Representation of the initial (UP) and final orientations in the 2D tilt plane (the points labelled a, b, c, d correspond to the respective panels). In general, starting from UP, the head may reach α=60° with any orientation γ (defined in **Suppl. Fig. 1**). These possible orientations are represented by a blue line and correspond to any tilt position that is exactly 60° from UP. This rule can be generalized as follows: *following a rotation of Axis IV by x°, the head may reach any tilt position that is exactly x° from its initial tilt, depending of its initial orientation in the horizontal plane*. **(f-g)** Illustration of Experiments 3-L,T: the 3D rotation protocol (f) is repeated after tilting the entire enclosure (g). **(h)** Responses of 3 hypothetical cells are shown in red, green and blue. *Top:* Hypothesis H_1_. We expect that neuronal tuning will appear re-organized (each hill of activity being transformed in a circular tuning curve based on the rule illustrated in panels a-e) in a Visual reference frame (compare middle vs. left column). In contrast, tuning will remain relatively unchanged when expressed in a Gravity reference frame (right vs. left column). Bottom: Hypothesis H_2_ with reverse properties.

**Supplemental Figure 15:**
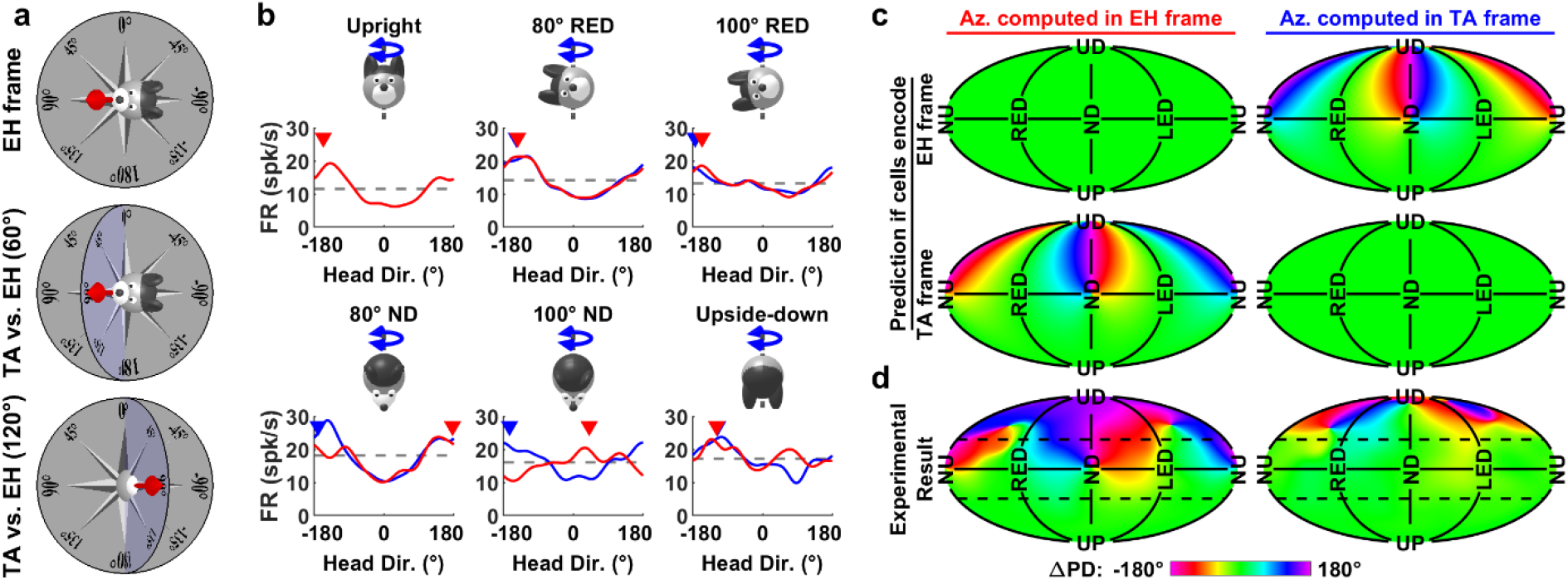
Azimuth tuning is spatially invariant when expressed in a Tilted (TA) frame. **(a)** Illustration of the EH and TA model (equivalent to the dual-axis rule; see Page et al., 2017; Laurens and Angelaki, 2018). In an EH frame (top panel), head direction is projected onto the EH plane (grey). In a TA frame, head direction is measured in a compass (blue) that is coplanar with the head horizontal plane and oriented such that the azimuth measured in the EH and TA frame coincide along the line of intersection of both planes (the 0-180° axis here). In other words, the TA frame is anchored to the allocentric reference frame along the earth-horizontal direction. It is defined by rotating a horizontal compass to align with head direction, instead of projecting head direction onto the horizontal plane (EH frame). In the example orientations shown here, the head pitches upward by 60° (middle panel) and 120° (bottom panel). In a TA frame, it faces 90° in both panels. When projected on the EH plane, its direction reverses from 90° to −90° when pitch angle exceeds 90°, as reported by Finkelstein et al. (2015). Note that, if the head is facing the 0° (or 180°) direction, it would be rolling instead of pitching, and azimuth reversal would not occur since these directions coincide in the EH and TA frames. As a general rule, TA is reversed relative to EH azimuth when tilt angle exceeds 90° in the pitch plane, but not in the roll plane. In intermediate tilt planes, the difference between EH and TA azimuth depends of tilt angle (see panel c). **(b)** Azimuth tuning curves of an example cell, extracted from the full 3D tuning curve measured in Experiment 3-L and computed in EH (red) or TA (blue) reference frames. Each curve represents the firing rate for all possible azimuths at a single tilt angle, which correspond to the positions attained by tilting the head to a given orientation and rotating around an earth-vertical axis (see **Suppl. Movie** 1). Note that the azimuth response of the example cell is modest since azimuth tuning is reduced when measured in a rotator (see **Suppl. Fig. 8a,b**). In an EH frame, the cell’s PD (−157°) was conserved for tilt orientations in the roll plane (−143° and −156° at 80° and 100° RED) but reversed abruptly for tilt angles larger than 90° in the pitch plane (from 177° to 41° at 80° and 100° ND). However, there is no such abrupt reversal when azimuth is computed in the TA frame. **(c)** Predicted change of the cell’s azimuth PD in tilted orientations (ΔPD, expressed relative to its PD when upright), displayed as a color map. For a given 3D head orientation, azimuth differs when computed in a EH or TA frame. The difference between both azimuths depends on the head’s orientation relative to gravity (see **Online Methods;** Page et al., 2017; Laurens and Angelaki, 2018). If cells encode azimuth in an EH frame, we expect their PD to be invariant across all head tilts (i.e. ΔPD≈0) when azimuth is expressed in the EH frame (upper left panel) but to vary with head tilt, and in particular to reverse when the head is pitched beyond 90° (i.e. between NU/ND and UD) when azimuth is expressed in TA frame (upper right panel). Reciprocally, if cells encode azimuth in a TA frame, we expect their PD to be invariant when expressed in a TA frame (lower right panel) but to vary when expressed in EH frame (lower left). **(d)** Average ΔPD across all azimuth-tuned cells significantly tuned to azimuth in Experiment 3-L (17 ADN, 7 RSC, 39 CIN). The azimuth PD varies when expressed in an EH frame but remains invariant (except close to UD) when expressed in a TA frame, in line with predictions of the TA model. Note that the PD becomes more variable close to UD, likely because azimuth tuning amplitude is near zero close to UD (see **Fig. 5g**), making data unreliable at this orientation.

**Supplemental Figure 16:**
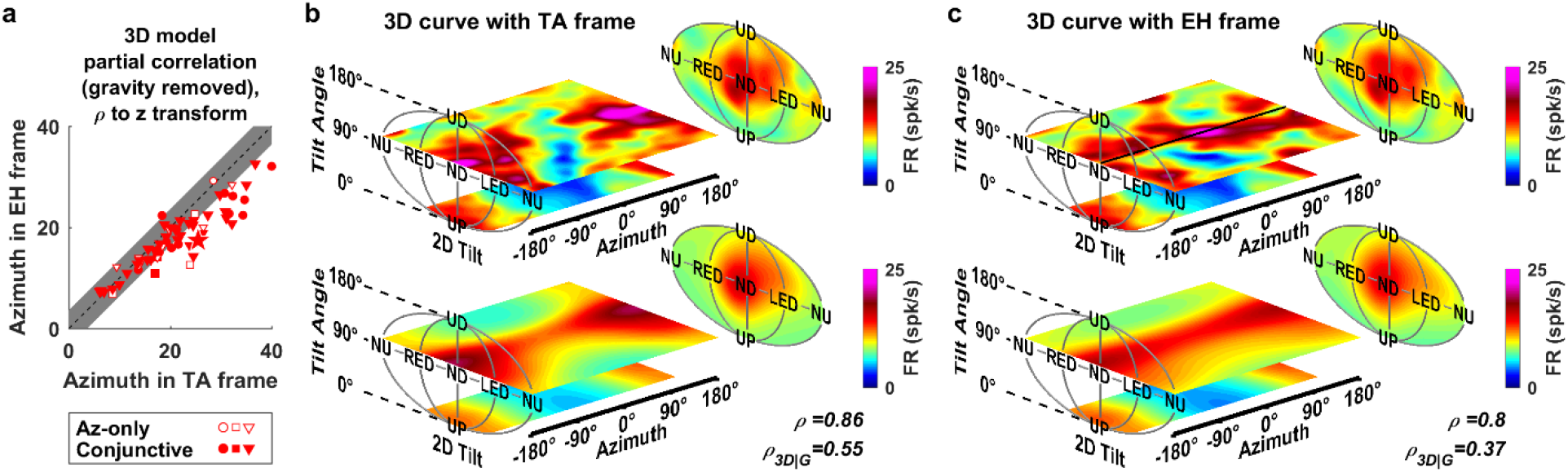
Comparison of 3D azimuth/tilt separable model fitting performed with azimuth in TA or EH frame. **(a)** Partial correlation of the model fits (shown as z-score), with the correlation attributable to gravity removed so that the partial correlation reflects how the model fits azimuth tuning in 3D. Data for all Az-tuned neurons that maintained their azimuth tuning in the rotator (Experiment 3-L) when the head is close to upright (<45° tilt; 17 ADN, 7 RSC, 39 CIN). Grey band: zone where partial correlations are not significantly different at p<0.01. Partial correlations were significantly higher when azimuth was expressed in a TA frame in 24/63 neurons, and significantly higher in a EH frame in only 1 ADN neuron (in this neuron, the difference between both frames was weak and vanished if only data for > 90° tilt was analyzed, indicating that it is likely a false positive). This analysis confirms that neuronal responses are more consistently expressed in a TA frame. The absence of significant difference in a large fraction of neurons (38/63) is explained by both the similarity between TA and EH frames at small tilt angles and the tendency of azimuth responses to decrease with tilt angle (see **Fig. 5f,g**). An alternative explanation, which would be that cells encode a mixture of EH and TA azimuth, may be rejected because the two frames are mutually exclusive. **(b,c)** 3D tuning curve of an example neuron computed in both frames (upper panels) and corresponding model fits (lower panels). This neuron was tuned to tilt, with a PD at α=100° tilt in ND orientation (γ=-5°), as well as azimuth with a PD at −175° when upright (lower planes in the 3D curve). TA and EH frames are identical near upright and, accordingly, tuning appears similar. Next, we examine tuning at a tilt angle of 100° (upper planes), where the TA and EH frames diverge sharply (as in **Suppl. Fig. 15a,c**). In a TA frame (b), the cell still exhibited a clear azimuth tuning with a similar PD (168°) as in upright. The 3D model (lower panel) captured the 3D curve by multiplying a tilt tuning centered on 100° ND with an azimuth tuning curve centered on 175°, leading to a total correlation of ρ=0.86 and a partial correlation of ρ_3D|G_=0.55. In contrast, azimuth tuning was largely distorted when expressed in a EH frame (upper plane on panel c). In ND orientation (marked by a black line), the cell’s response reversed and peaked at an azimuth of 18° (magenta). In contrast, it shifted back to ±180° on either side of the line, i.e. when head orientation neared RED and LED. This pattern, where ***azimuth tuning reverses in ND but not RED or LED*, corresponds to the reversal of TA azimuth relative to EH azimuth (**Suppl Fig. 15a,c**) and is *expected if azimuth is encoded in a TA frame***. Therefore, the cell’s azimuth PD was not invariant relative to head tilt when expressed in an EH frame. Since this violates the assumption of the 3D model, the correlation decreased to ρ=0.8 and ρ_3D|G_=0.37 (note that the correlation didn’t decrease to zero since the model could still fit azimuth tuning at low tilt angles).

**Supplemental Figure 17:**
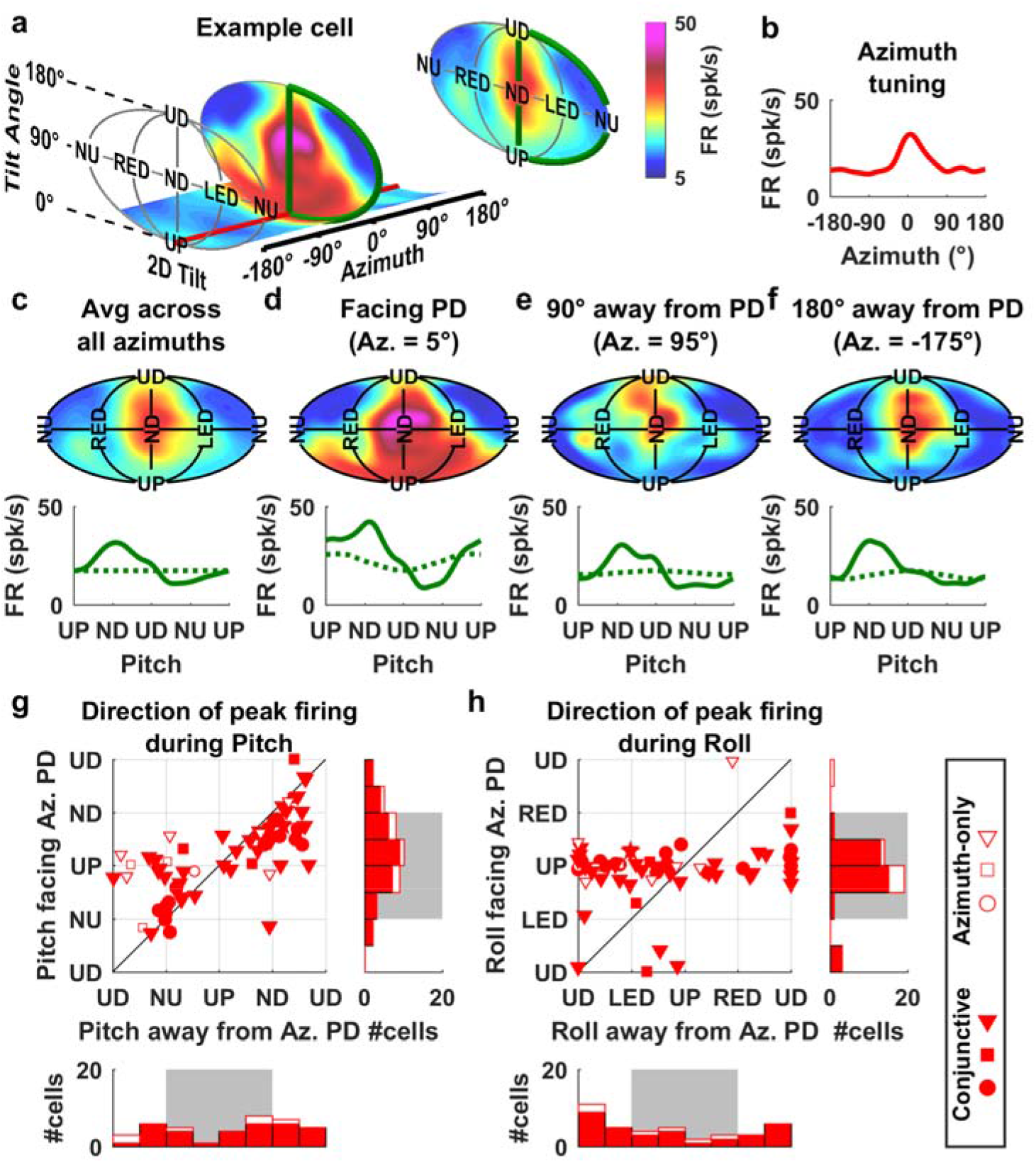
Possible bias when measuring pitch and roll tuning in azimuth-tuned cells. (see also Laurens and Angelaki 2019). A recent study in rat ADN (Shinder and Taube, 2019) failed to identify tilt responses in a sample of 24 azimuth-tuned HD cells. In that study, mice were positioned upright, facing the azimuth PD, and rotated in pitch and/or roll. The authors observed that most cells fired more in tilt positions near upright and concluded on that account that there is “limited evidence that cells contained conjunctive firing with pitch or roll position” (sic). Here we demonstrate that the experimental protocol used by Shinder and Taube (2019), where mice were tilted while facing the cell’s PD, tends to conceal pitch/roll tuning, because it is superimposed on a strong azimuth tuning, whose strength is reduced as a function of head tilt (**Fig. 5f,g**). **(a)** 3D tuning curve of an example conjunctive cell (measured during Experiment 3-L; same cell as in **Fig. 6c** and **Suppl. Fig. 13**). When averaged across all azimuths (rightmost plane), the cell is tuned to tilt with a PD in ND. The cell is also tuned to azimuth, with a PD at 5°. A vertical section (i.e. firing rate for all tilt position at a given azimuth) of the tuning curve is shown at the azimuth PD. When exclusively tested during pitch in this plane (green line; as Shinder and Taube, 2019, did), the cell’s tilt modulation is much broader and the cell’s firing at ND is barely above its firing when the animal is upright (red). **(b)** Average azimuth tuning curve when upright, peaking at 5°. **(c)** Analysis in this study: Upper panel: Tilt tuning curves averaged across all azimuth angles (as in panel a). Lower panel: Firing rate measured along a pitch trajectory; Solid green curves: actual data; Dashed green curves: simulated data (see below). **(d)** Experiment by Shinder and Taube (2019): Tilt tuning was tested when animals faced the azimuth PD. Note that firing is elevated in the vicinity of upright. Even though the firing is largest in ND, the preferred pitch direction, evaluated by fitting a von Mises function, is biased towards upright (67° ND tilt in d versus 108° ND tilt in c). **(e,f)** Tilt tuning tested 90° (e) or 180° (f) away from the cells’ azimuth PD. Firing measured during pitch rotation away from the PD is similar as the average curve in **(c)**. We use the 3D model fit to demonstrate that the curve in (d) is affected by azimuth tuning. We fit the cell’s 3D tuning curve, then alter the model’s parameter to eliminate tilt tuning (by setting A to 0 and FR_0_ to the cell’s average firing in FR_Ti_(α,γ); see **Methods**). Next we simulate the “pitch tuning curve” (dashed green curves) that is now influenced entirely by its azimuth tuning. The resulting curve peaks in UP orientation in (d), but is flat in other panels. This indicates that azimuth tuning affects the cell’s response when facing the azimuth PD (d), such that it biases the firing rate towards upright (by interacting multiplicatively) but has little effect when facing away from the azimuth PD (e,f) or when data are averaged across all azimuths (c). We note that ***the green curve in (d) resembles most example pitch or roll tuning curves shown by Shinder and Taube (2019)***. Based on these simulations, we predict that, had the authors analyzed individual pitch/roll tuning curves recorded when the mouse faced away from the cell’s PD (e, f), they would have seen tilt tuning with preferred tilt away from upright. **(g,h)** Same analysis, at the population level. We simulated pitch/roll rotations for all azimuth-tuned cells that were also azimuth tuned in the rotator (n=63; 53 conjunctive and 10 azimuth-only cells). Top: Scatter plot showing how pitch rotations while facing the cell’s azimuth PD can bias conclusions. Peak responses (by fitting von Mises functions) to pitch and roll rotations when facing the azimuth PD (ordinate; as in Shinder and Taube, 2019) and when facing away from the azimuth PD (abscissa). Bottom and right: Marginal distributions are shown as histograms. When pitching while facing the azimuth PD (panel g), most conjunctive cells fire preferentially close to upright right-side histogram, grey zone, red bars (41/53, 78%, p<10^−4^), similar to the example cell in a-f. When adding azimuth-only cells (open symbols/bars), the proportion of cells firing preferentially close to upright is maintained at 49/63 (77%). The bias is even more drastic in roll (panel h; because tilt tuning is weaker in roll), with the 49/53 conjunctive cells firing preferentially around upright (lower histogram, grey zone, red). In contrast, when pitching or rolling away from the azimuth PD (g,h; abscissae), half of the conjunctive cells (solid red symbols/bars, 24/53 in g, 19/53 in not significantly different from 50%, p=0.6/0.05 respectively) fire preferentially close to upright (grey band in the marginal distribution) and the other half fire preferentially closer to UD. Thus, while recording from ‘our’ neurons. ***if we had done the experiment (pitch/roll when animal faced azimuth PD in a small sample of cells) and analyses as in Shinder and Taube (2019), we would likely not have been able to identify tilt tuning***. We conclude that our dataset and quantitative analyses predict that, even though the PD of tilt tuning is distributed uniformly between upright and inverted orientation (**Fig. 3a**), conjunctive cells would appear to respond preferentially in upright orientation when recorded and analyzed as in (Shinder and Taube, 2019). The results published in our study are therefore entirely compatible with those described by Shinder and Taube (2019). The conclusions are opposite because the systematic scanning of 3D orientation (rather than a limited subset) allowed us to reveal tilt tuning, that was concealed by azimuth tuning in the Shinder and Taube (2019) study.

**Supplemental Figure 18:**
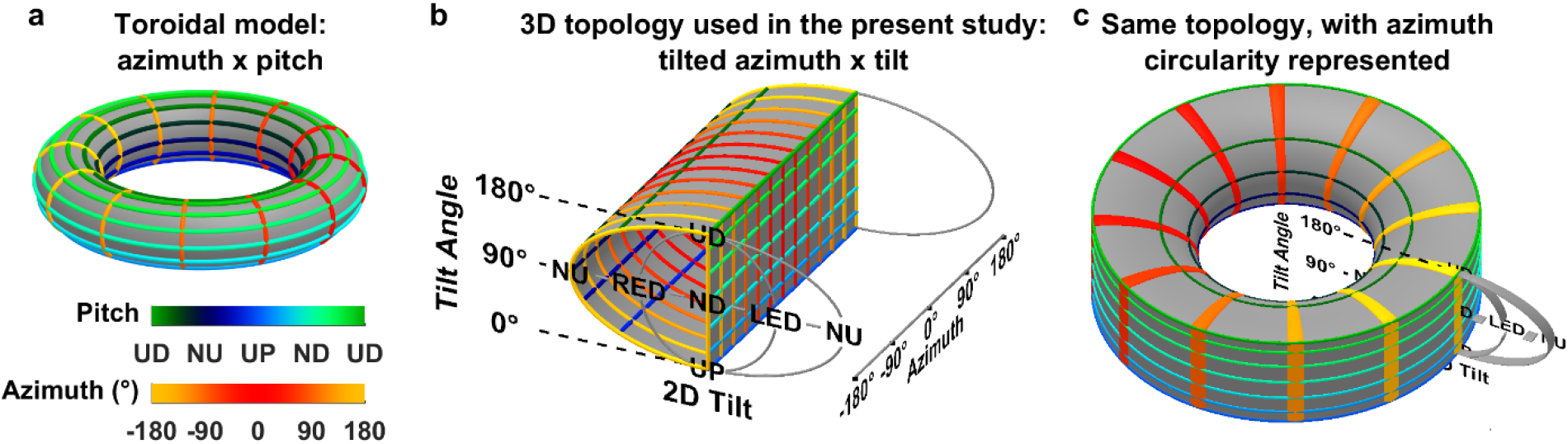
Comparison between the separable, multiplicative model of Fig. 6 and toroid topology (Finkelstein et al. 2015). **(a)** The toroid model is restricted to tilt movements in pitch, and assumes that azimuth and pitch are independent, i.e. pitch movements don’t change azimuth. Combination of pitch and azimuth can be represented on the surface of a torus. ‘Iso-pitch’ lines (green/blue/black color code), that correspond to one pitch orientation and all possible azimuths, form horizontal circles. ‘Iso-azimuth’ lines (yellow/red color scale), that correspond to one azimuth angle and all possible pitch tilts form vertical lines. **(b)** Representation of the same iso-pitch and iso-azimuth lines in the 3D topology used in this study, when azimuth is expressed in a TA frame. Each isoazimuth line forms a ‘D-shapeď curve that passes through UP, NU, UD and ND orientation, and each iso-pitch line forms a horizontal line. **(c)** When the diagram in (b) is ‘looped’ upon itself to account for the circularity of azimuth (note that this representation was not used outside of this figure because it distorts volumes), the surface formed by iso-azimuth and iso-pitch lines adopts a toroidal topology identical to (a). Thus, the 3D model used here is equivalent to the toroidal topology in (Finkelstein) if (1) tilt is restricted to the pitch plane and (2) azimuth is expressed in the TA frame.

**Supplemental Figure 19:**
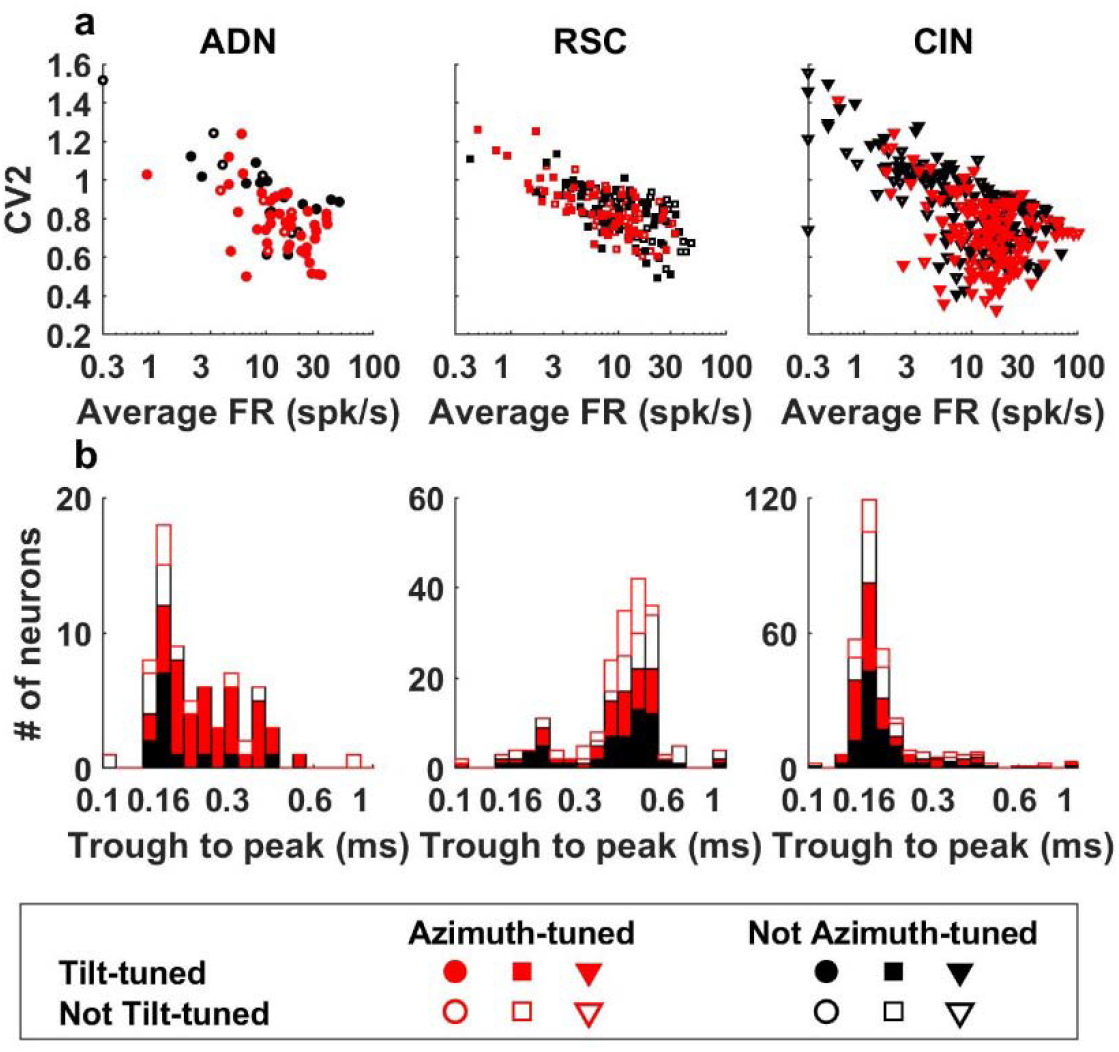
Spike waveform and firing properties in sampled areas. **(a)** Scatter plot of CV2 vs. average firing rate during freely moving in the arena. Median firing rate: 14 spk/s in ADN, 10 spk/s in RSC, 13 spk/s in CIN. Median CV2: 0.83 in ADN, 0.84 in RSC, 0.79 in CIN. Firing rate varied significantly across areas (Kruskal-Wallis non-parametric ANOVA, p = 0.006), but CV2 was similar (p=0.04). **(b)** Trough to peak duration of action potentials. The median duration in ADN and CIN (0.2 ms and 0.17 ms, respectively) was significantly smaller (Kruskal-Wallis non-parametric ANOVA, p<10^−10^) than in RSC (0.44 ms). The CIN is a fiber bundle and neuronal activity recorded therein is therefore expected to consist of axonal spikes, that can be recorded by tetrodes (Robbin et al. 2013) and typically exhibit small duration (Barry, 2015). Note that most units recorded in the ADN also had short-duration spikes. To our knowledge, no studies has reported the spike duration of HD cells in the ADN. Investigating the physiology of neuronal activity underlying ADN HD cells would be an interesting topic for future studies.

